# Glutaminase as a metabolic target of choice to counter acquired resistance to Palbociclib by colorectal cancer cells

**DOI:** 10.1101/2024.01.04.574237

**Authors:** Míriam Tarrado-Castellarnau, Carles Foguet, Josep Tarragó-Celada, Marc Palobart, Claudia Hernández-Carro, Jordi Perarnau, Erika Zodda, Ibrahim H. Polat, Silvia Marin, Alejandro Suarez-Bonnet, Juan José Lozano, Mariia Yuneva, Timothy M. Thomson, Marta Cascante

**Affiliations:** Department of Biochemistry and Molecular Biomedicine and Institute of Biomedicine (IBUB), Universitat de Barcelona, Barcelona, Spain; CIBER of Hepatic and Digestive Diseases (CIBEREHD), Institute of Health Carlos III (ISCIII), Madrid, Spain; British Heart Foundation Cardiovascular Epidemiology Unit and Victor Phillip Dahdaleh Heart and Lung Research Institute, University of Cambridge, Cambridge, UK; Oncogenes and Tumour Metabolism Laboratory, The Francis Crick Institute, London, UK; Experimental Histopathology, The Francis Crick Institute, London, UK; Department of Pathobiology and Population Sciences, The Royal Veterinary College, Hatfield, UK; Bioinformatics Platform, Centro de Investigación Biomédica en Red Enfermedades Hepáticas y Digestivas (CIBEREHD), Barcelona, Spain; Institute for Molecular Biology (IBMB-CSIC), Barcelona, Spain; Instituto de Investigaciones de la Altura, Universidad Peruana Cayetano Heredia, Lima, Peru; Instituto de Investigaciones Científicas y Servicio de Alta Tecnología (INDICASAT AIP), Panama City, Panama

## Abstract

Several mechanisms of resistance of cancer cells to cyclin-dependent kinase inhibitors (CDKi) have been identified, including the upregulation of metabolic regulators such as glutaminase. However, whether such mechanisms and targets are optimal has not been determined. Here, we have systematically analyzed metabolic reprogramming in colorectal cancer cells exposed to Palbociclib, a CDKi selectively targeting CDK4/6, or Telaglenestat, a selective glutaminase inhibitor. Through multiple approaches, we show that Palbociclib and Telaglenestat elicit complementary metabolic responses and are thus uniquely suited to counter the metabolic reprogramming induced by the reciprocal drug. As such, while Palbociclib induced reduced tumor growth *in vivo*, and Telaglenestat did not show a significant effect, the drug combination displayed a strong synergistic effect on tumor growth. Likewise, initial responses to Palbociclib were followed by signs of adaptation and resistance, which were prevented by combining Palbociclib with Telaglenestat. In conclusion, combination with Telaglenestat optimally forestalls acquired resistance to Palbociclib in cancer cells.

## Introduction

Cell cycle control is frequently dysregulated in cancer cells leading to unscheduled proliferation (Diaz-Moralli et al., 2013). Cyclin-dependent kinases CDK4 and CDK6 (CDK4/6) are promising targets in cancer therapy since their overexpression and dysregulation are implicated in a wide range of human cancers (Goel et al., 2022; Sherr et al., 2016). Three selective pharmacologic CDK4/6 inhibitors have received approval from the US Food and Drug Administration (FDA) and the European Medicines Agency (EMA) for the treatment of hormone receptor (HR)-positive, human epidermal growth factor receptor 2 (HER2)-negative locally advanced or metastatic breast cancer (Groenland et al., 2020). Among these, Palbociclib (PD0332991) was the first-in-class drug to show promising results in phase III studies (Beaver et al., 2015; Marra and Curigliano, 2019; O’Leary et al., 2016). More recently, Palbociclib and other CDK4/6 inhibitors (CDK4/6i), in a variety of combinations, have shown efficacy in other cancer types, including colorectal carcinoma (Bride et al., 2022; Coffman et al., 2022; de Kouchkovsky et al., 2022; Gleason et al., 2024; Kase et al., 2024; Lee et al., 2023; Lee et al., 2016; Lelliott et al., 2021; Mao et al., 2021; Martin-Broto et al., 2023; Movva et al., 2024; Ngamphaiboon et al., 2024; Pek et al., 2017; Pesch et al., 2022; Raetz et al., 2023; Selby et al., 2023; Sorokin et al., 2022), particularly in tumors bearing genetic or epigenetic alterations underlying an upregulated G1-S cell cycle transition (Mao et al., 2021; Pesch et al., 2022). In addition to a direct effect on cancer cell survival, CDK4/6i have been shown to promote anti-tumor immune responses (Lelliott et al., 2021; Petroni et al., 2020). However, as single agents, CDK4/6i have shown limited efficacy outside of ER+ breast cancers (Clark et al., 2023; Zeverijn et al., 2023) and, like with most targeted therapies, tumor cells eventually acquire resistance to CDK4/6 inhibition (Álvarez-Fernández and Malumbres, 2020; McCartney et al., 2019; Park et al., 2024; Wingate and Keyomarsi, 2023). Experimentally tested mechanisms of CDK4/6i resistance include gain-of-function changes impacting cell-cycle transitions (Freeman-Cook et al., 2021; Gomatou et al., 2021; Li et al., 2022; Zikry et al., 2024), MYC dependency (Freeman-Cook et al., 2021; Ma et al., 2024; Park et al., 2024; Tarrado-Castellarnau et al., 2017), metabolic reprogramming (Santiappillai et al., 2021; Tarrado-Castellarnau et al., 2017; Zhou et al., 2023), or other signaling and regulatory pathway adaptations (Goodwin et al., 2023; Jagirdar et al., 2024; O’Brien et al., 2020; Park et al., 2024; Rodriguez et al., 2023).

Our previous work accurately characterized the metabolic reprogramming associated with CDK4/6 pharmacological inhibition or genetic depletion, uncovering the upregulation of MYC, glutaminolysis, and mTOR signaling as major adaptations of colorectal cancer cells to restore their fitness in response to such perturbations (Tarrado-Castellarnau et al., 2017). Our results demonstrated that these adaptations cancer cells that survive CDK4/6 inhibition highly sensitive to MYC, glutaminase, or mTOR inhibitors and hypoxic conditions. As such, we validated the combined inhibition of CDK4/6 and glutaminase (GLS1) as an effective strategy to forestall resistance to CDK4/6 therapy with strong synergistic and selective antiproliferative effects on tumor cells *in vitro*.

Herein, we have analyzed the metabolic reprogramming associated with the inhibition of CDK4/6 or glutaminase (GLS1), and their combination, both *in vitro* and *in vivo*, in a mouse xenograft model of HCT116 human colorectal cancer cells. Using genome-scale metabolic modeling, we have integrated transcriptomics and metabolomics data to reveal, systematically and in an unbiased fashion, the most relevant up- and down-regulated metabolic pathways in colorectal cancer cells in response to single or combined *in vivo* therapies. Our results establish that upregulation of glutaminolysis is the most prominent metabolic mechanism explaining resistance to Palbociclib. As such, combining Palbociclib with the selective GLS1 inhibitor, Telaglenestat (CB-839), counteracts the adaptive response of tumor cells to Palbociclib both *in vitro* and *in vivo,* thus averting the acquisition of pharmacological resistance. Remarkably, we have found that residual cancer cells from tumors treated *in vivo* with the combined therapy remain sensitive to the dual treatment *ex vivo*, long after chemotherapeutic exposure, and exhibit a downregulated metabolism. Together, our results validate the combination of Palbociclib and Telaglenestat as an effective strategy to prevent pharmacologic resistance and potentiate the antiproliferative effects of single-agent treatments.

## Results

### Metabolic adaptations of HCT116 cells to prolonged treatment with Palbociclib and Telaglenestat

Our previous investigations revealed that the combined therapy with CDK4/6 and glutaminase inhibitors presented strong antiproliferative synergies in colon cancer cells with no significant cytotoxic effects on non-malignant cells (Tarrado-Castellarnau et al., 2017), predicting for this combination a promising therapeutic use in cancer treatment. As such, we first characterized *in vitro* the antiproliferative effects of Telaglenestat (CB-839) (Supplementary Figure 1A), alone and in combination with Palbociclib (PD0332991), since it is a selective glutaminase inhibitor of both splice variants of glutaminase (KGA and GAC) with good oral bioavailability (Gross et al., 2014; Harding et al., 2021), currently in phase II clinical trials (NCT04265534). In contrast, the classical glutaminase inhibitor, BPTES, has poor metabolic stability and low solubility, limiting its potential for *in vivo* assays and requiring further clinical development (Shukla et al., 2012). The combination of Palbociclib and Telaglenestat presented strong synergistic antiproliferative effects over a wide dose range (CI < 0.3) (Figure 1A and Supplementary Figure 1B) on HCT116 colorectal cancer cells, as well as a reduction in anchorage-independent growth (Figure 1B) and clonogenic potential (Figure 1C). We also observed a synergistic effect on cell cycle arrest at the G1 phase for the combined condition (Figure 1D). Unexpectedly, while Palbociclib imposed a pronounced antiproliferative effect on HCT116 cells (Figure 1E,F and Supplementary Figure 1C), the cell doubling time was only marginally increased (Figure 1G), suggesting an early inhibition of cell division with a subsequent adaptation to this drug. Glutaminase inhibition alone had no impact on cell proliferation or duplication time. In contrast, the combined treatment caused a significant rise in cell doubling time compared to all groups and, consequently, a greater inhibitory effect on cell proliferation (Figure 1E-G and Supplementary Figure 1C). To control for cell type-specific biases, since HCT116 cells exhibit microsatellite instability (MSI) (Boland and Goel, 2010), we examined the synergistic antitumor effects of the combined treatment in microsatellite stable (MSS) colon cancer cell lines from different Consensus Molecular Subtypes (CMS) (Sveen et al., 2018), including SW403 (CMS2), HT29 (CMS3), and SW620 (CMS4) cell lines. Our findings demonstrated that the antiproliferative synergistic effects of the combined treatment were independent of the CMS and microsatellite stability status (Supplementary Figure 1D-F), since the combination of Palbociclib and Telaglenestat synergistically inhibited the proliferation of all cell lines at all concentrations tested, suggesting that this combination may be effective across a wide range of CRC molecular profiles. Collectively, these results corroborate that glutaminase inhibition synergizes with Palbociclib to impair colon cancer cell proliferation *in vitro*.

**Figure 1.**
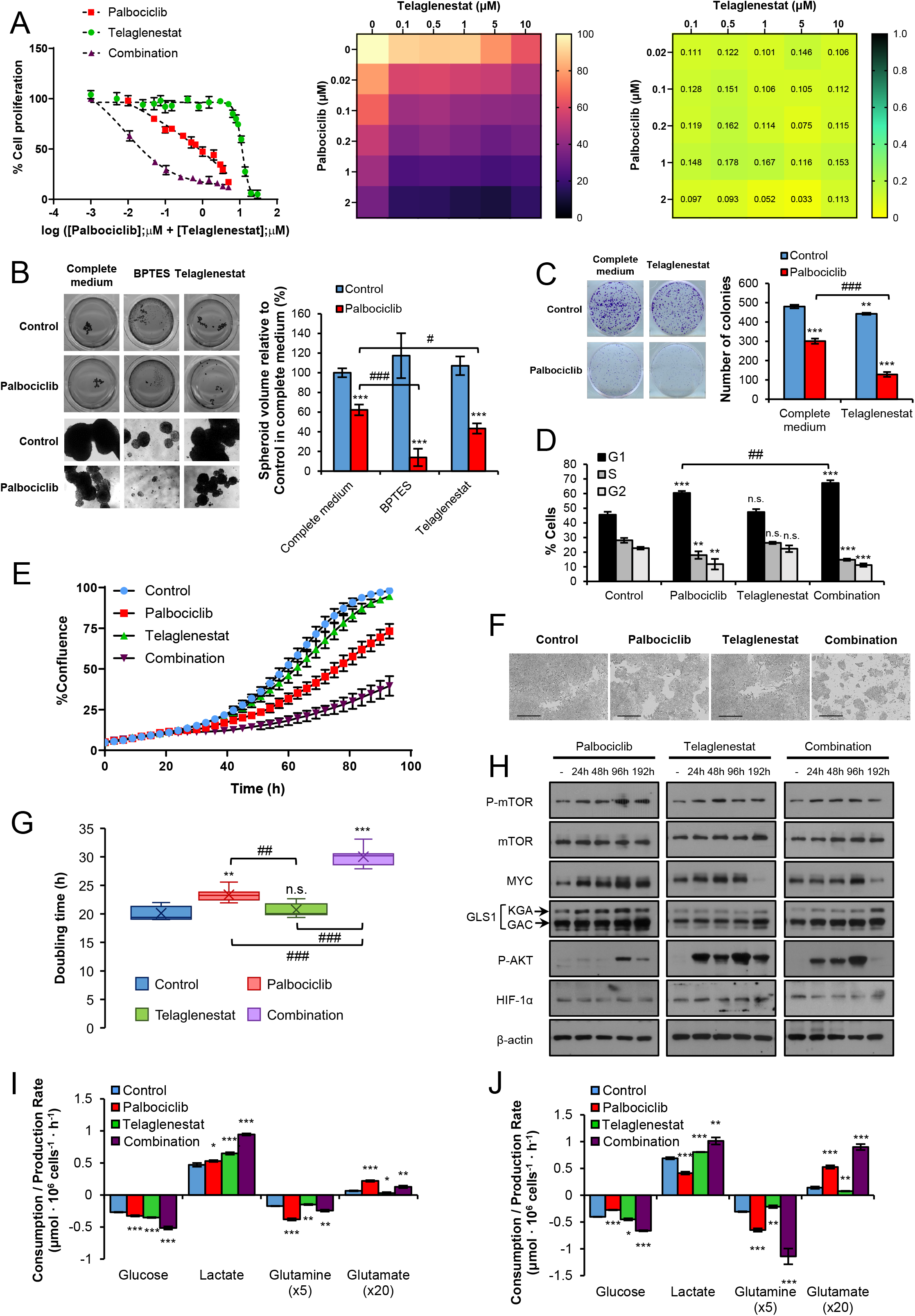
The antiproliferative effects of Palbociclib and Telaglenestat combination are synergistic and time-dependent. A. Cell proliferation assay. HCT116 cells were treated with increasing concentrations of Palbociclib, Telaglenestat, or their combination for 96 h, and cell quantification was assessed by Hoechst 33342 staining. Left, concentration-response curves generated by data fitting with a four-parameter equation. Center, dose-response cell proliferation matrix. Results are shown as the percentage of proliferation relative to untreated cells (mean ± SD for n=6). Right, dose-response synergy matrix. The combination index (CI) values were obtained with CompuSyn software (ComboSyn, Inc., Paramus, NJ, USA), revealing a strong synergy (CI<0.3) at each dose combination tested. B. Spheroid formation assay. Left, scan (top) and phase-contrast microscope (60x, bottom) images of HCT116 spheroids grown on anchorage-independent conditions and treated for 10 days with Palbociclib, the glutaminase inhibitors BPTES and Telaglenestat, or their combination. Right, quantification of total spheroid volume. Spheroids were scored by image acquisition, followed by spheroid area and volume quantification with ImageJ. Results are represented as the percentage of total spheroid volume relative to untreated cells (mean ± SD of n=4). C. Colony-formation assay. Left, images of HCT116 cells cultured in the absence or presence of 2 µM Palbociclib, 5 µM Telaglenestat, or their combination for 10 days. Right, quantification of the total number of colonies scored by image acquisition with ImageJ. D. Cell cycle distribution in HCT116 cells treated with Palbociclib, Telaglenestat, or their combination for 96 h and determined by flow cytometry. E. Cell proliferation curves for HCT116 cells treated with Palbociclib, Telaglenestat, or their combination for 96 h. Confluence was monitored in an IncuCyte^®^ S3 (Sartorius) live-cell analysis system. Results are shown as mean ± SD with n=6. F. Representative IncuCyte^®^ S3 live-cell images in phase-contrast of HCT116 cells treated with Palbociclib, Telaglenestat, or their combination at 96 h. Cell confluence was monitored every 3 h. Scale bar = 400 μm. G. Cell doubling time of HCT116 cells treated with Palbociclib, Telaglenestat, or their combination. Shown values are the mean ± SD of six independent experiments with six replicates each. H. Effects of Palbociclib, Telaglenestat, or their combination on signaling pathways with time. P-mTOR, mTOR, MYC, GLS1, P-AKT, and HIF 1α protein levels were determined by Western blotting of total protein fractions using β-actin as a loading control. I, J. Comparative extracellular metabolic fluxes for HCT116 cells treated with Palbociclib, Telaglenestat, or their combination for (H) 96 h and (I) 192 h. Glucose and glutamine consumption and lactate and glutamate production rates were obtained after 24 h of incubation with fresh media and normalized to cell number. Data information: Concentrations used were 2 µM for palbociclib, 5 µM for Telaglenestat and 10 µM for BPTES. Shown values are mean ± SD for n=3 (except otherwise indicated). Significance was determined by one-way ANOVA and two-tailed independent sample Student’s t-tests. Statistically significant differences between treated and control cells are indicated as *P* < 0.05 (*), *P* < 0.01 (**), and *P* < 0.001 (***), while differences between treatments are shown as *P* < 0.05 (#), *P* < 0.01 (##), and *P* < 0.001 (###).

To determine whether the cellular response triggered by the inhibitors was exposure-time dependent, we treated HCT116 cells with Palbociclib, Telaglenestat, and their combination for 24, 48, 96, and 192 h. By analyzing protein levels of central metabolism regulators, we observed that Palbociclib caused an activation of the mTOR, MYC, and AKT signaling pathways, which reached peak levels at 96 h of treatment (Figure 1H). On the other hand, Telaglenestat induced an earlier activation of these pathways, as well as an upregulation of HIF-1α, while glutaminase protein levels were only increased at 192 h of treatment. Remarkably, the levels of MYC and P-AKT substantially decreased at 192 h in all cases, revealing a metabolic reprogramming dependent on time of drug exposure. We also observed that the drug combination prevented an early activation of MYC compared to either treatment alone.

Next, we assessed the cellular consumption and production rates of the major carbon sources at 96 and 192 h of treatment. In line with our previous results (Tarrado-Castellarnau et al., 2017), cells subjected to a 96 h incubation with Palbociclib displayed enhanced rates of glucose and glutamine consumption and of lactate and glutamate production (Figure 1I), which is also in agreement with the reported activation of mTOR, MYC, and AKT signaling pathways and increased protein levels of glutaminase (Figure 1H). However, after 192 h of treatment, the metabolic reprogramming of Palbociclib-treated cells shifted to a reduced glycolytic profile, while remaining highly glutaminolytic (Figure 1J). On the other hand, Telaglenestat partially inhibited glutaminolysis and increased glucose consumption and lactate production, both at 96 and 192 h. Interestingly, the metabolic adaptation triggered by the combined treatment displayed the greatest increase in glucose uptake and lactate release at both incubation times. The combination of inhibitors also prompted an increment in the glutaminolytic flux at 96 h compared to control and Telaglenestat-treated conditions, significantly increasing after eight days of treatment and in accordance with the upregulation of GLS1 protein levels reported at 192 h (Figure 1H).

### Palbociclib and Telaglenestat combined therapy presents synergistic antiproliferative effects *in vivo*

To validate our *in vitro* results, we investigated the effects of the combined administration of Palbociclib and Telaglenestat on the tumor growth of HCT116 cells xenografted into non-obese diabetic (NOD) severe combined immunodeficiency disease (SCID) mice. One week after HCT116 subcutaneous injection, mice were treated with either vehicle (Control), Palbociclib (100 mg/kg/day p.o.), Telaglenestat (150 mg/kg/day p.o.), or Palbociclib + Telaglenestat (100 mg/kg/day p.o. + 150 mg/kg/day p.o.) for 23 days (Supplementary Figure 2A). Previous studies determined that these doses were non-toxic (Fry et al., 2004; Gross et al., 2014) and mice tolerated all treatments without significant body weight loss (Supplementary Figure 2B-C). Of note, animals gavaged with the combined treatment gained less weight than the other three groups (Supplementary Figure 2D-E). Remarkably, while Palbociclib treatment alone induced a significant reduction in tumor volume compared to the control group (Figure 2A-C), the tumor growth rate of both groups was equivalent (Figure 2D-E). As a result, mice treated with Palbociclib alone did not display tumor growth inhibition (%TGI) compared to the control condition (Figure 2F). These results suggest that inhibition of CDK4/6 has only an early impact on tumor cell proliferation and that surviving cells can overcome this inhibitory effect and restore their growth. On the other hand, treatment with Telaglenestat alone failed to achieve a statistically significant reduction of tumor volume (Figure 2A-C).

**Figure 2.**
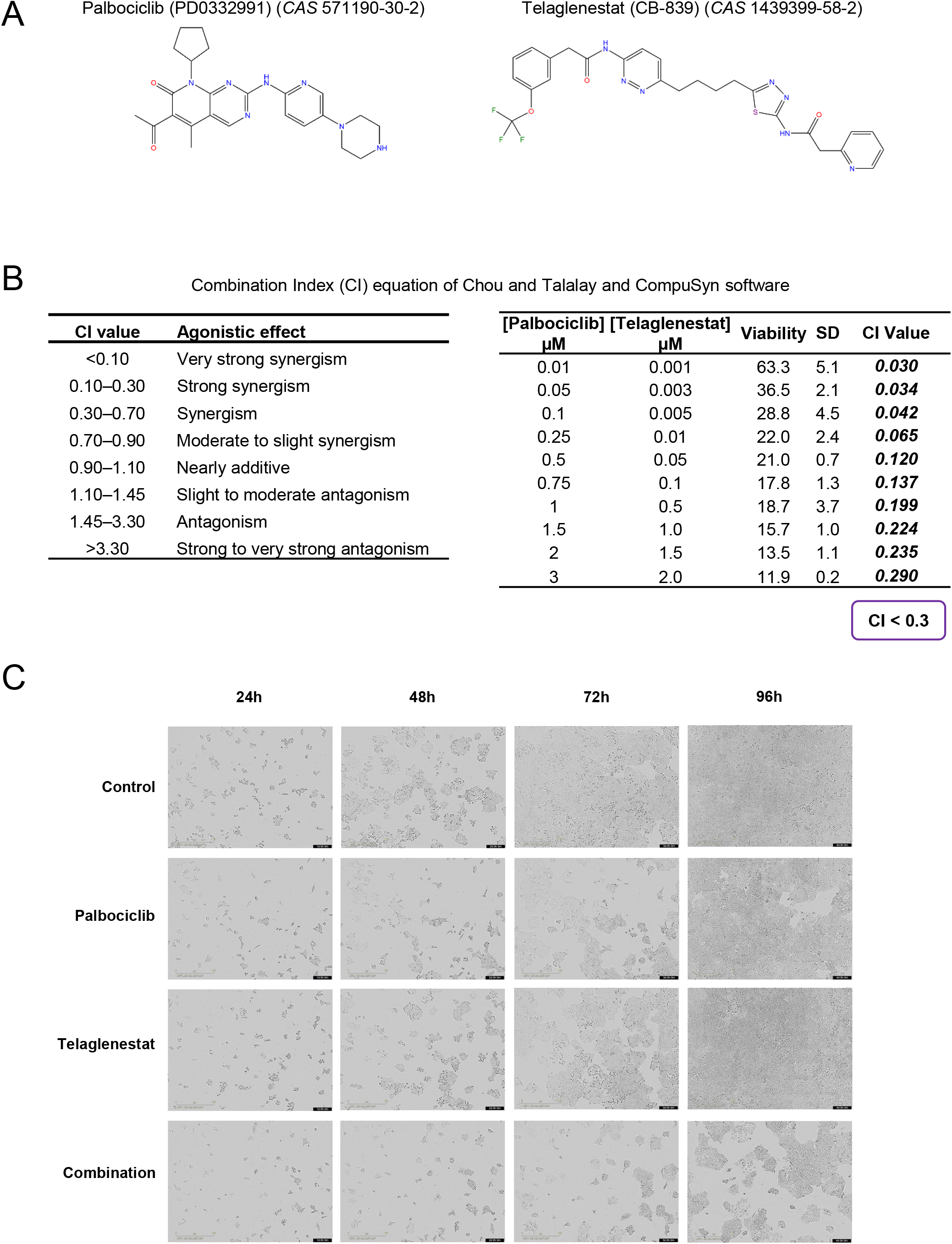
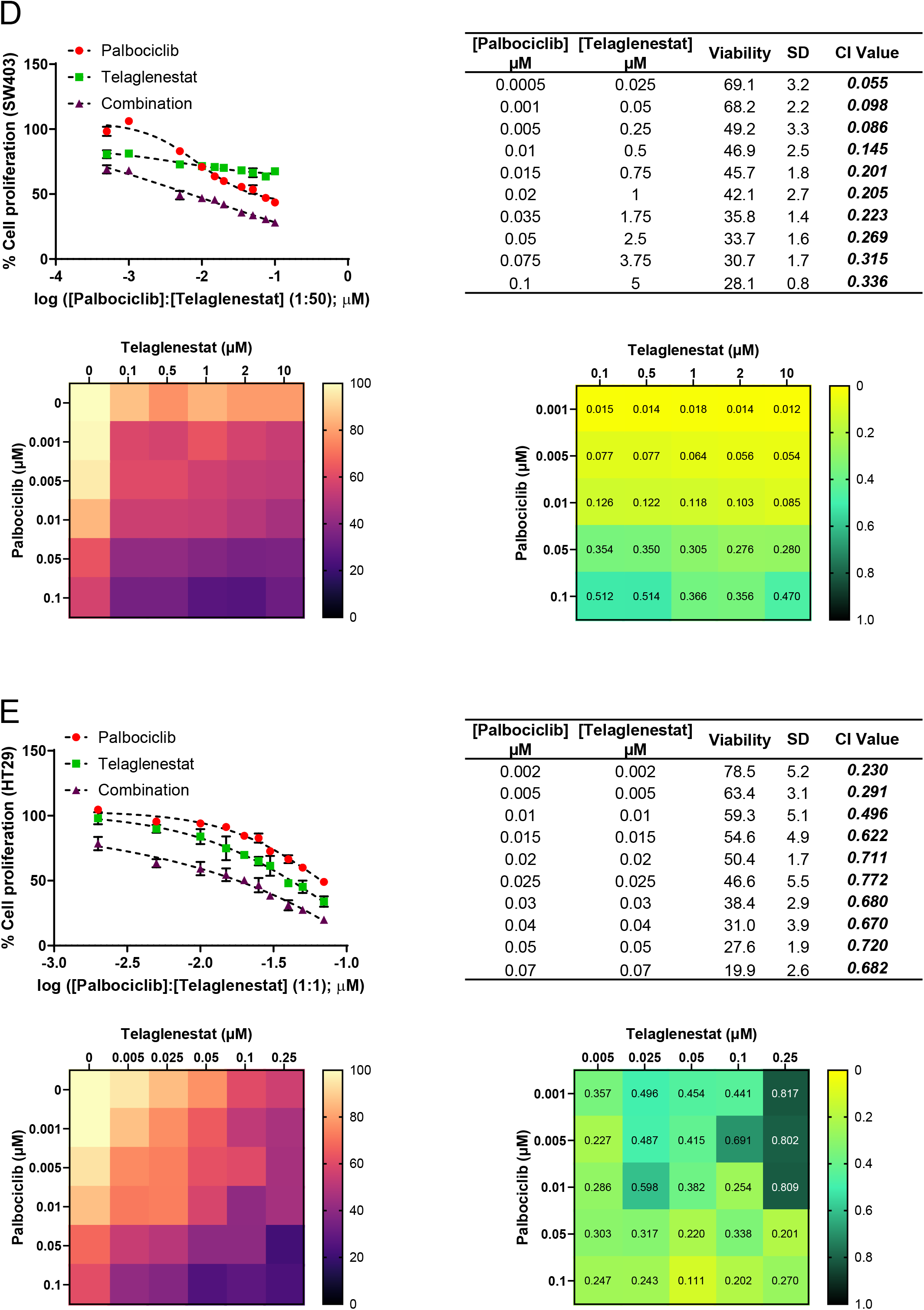
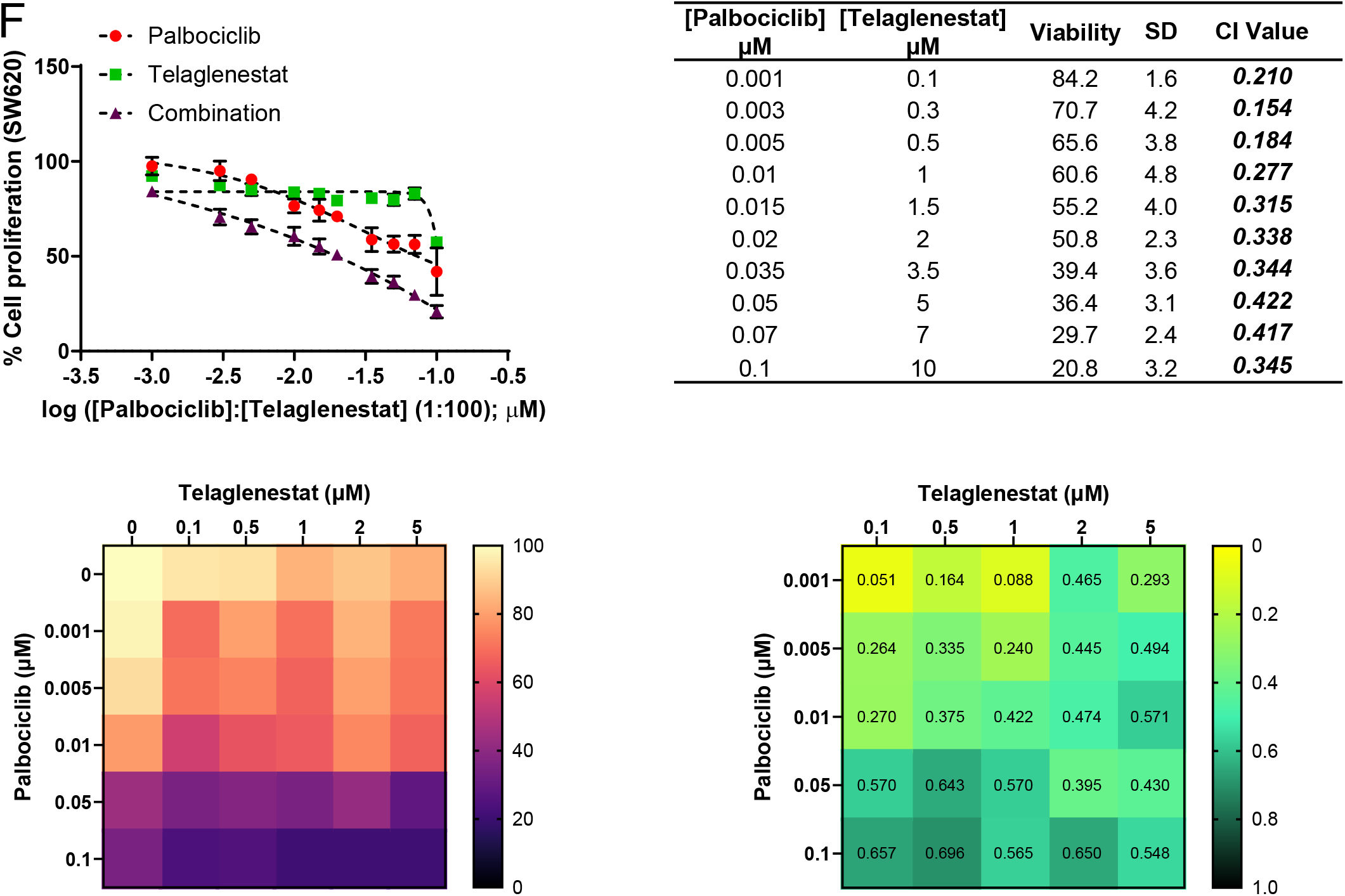
Palbociclib and Telaglenestat combined therapy significantly suppresses tumor growth *in vivo*. A. Growth curve of HCT116 xenografts. After one week of tumor establishment, NOD-SCID mice were treated daily for 23 days with vehicle (Control), Palbociclib, Telaglenestat, or their combination, and tumor volume was measured with a Vernier caliper at the indicated days using the modified ellipsoid volume formula: Tumor volume = (length × width^2^) × π/6 (n=12 per group). Data are presented as mean ± SEM. B. Final volume measured at necropsy (n=12 per group). Data are represented in a box and whiskers plot, with the whiskers representing the minimum and maximum values, all data points shown, and the median indicated. C. Representative *ex vivo* images of solid tumors from each treatment group excised from NOD-SCID mice on Day 31 after cell implantation. Scale bar = 1 cm. D. Tumor volume fold change from Day 18 (normalized to volume on Day 15) in HCT116 subcutaneous colorectal cancer xenografts (n=12 per group). Data are shown as mean ± SEM. E. Tumor volume fold change at Day 31 normalized to volume on Day 15 (n=12 per group). Data are represented in a box and whiskers plot, with the whiskers representing the minimum and maximum values, all data points shown, and the median indicated. F. Bioluminescence intensity relative to the first measurement time point of photon radiance (Day 8) in luciferase-expressing HCT116 xenografts (n=12 per group). Data are shown as mean ± SEM. G. Tumor growth inhibition (TGI) rate to compare the anti-tumor efficacy of the treatments from Day 18 (n=12 per group) calculated as TGI = (1-(V_(T,t)_/V_(T,0)_)/(V_(C,t)_/V_(C,0)_)) x 100, where time 0 is the first measurement of tumor volume on Day 15. Data are presented as mean ± SEM. H. Representative histopathology images of human colon carcinoma xenografts generated from HCT116 cells; magnification x200, scale bar = 50 μm. Top, Hematoxylin and Eosin (H & E) staining and mitotic figures quantification (mitotic count) expressed as the number of mitoses per 10 high-power fields (HPF), with HPF = 400x overall magnification. Middle, KI67 immunohistochemical staining and quantification of the percentage of KI67 positive cells. Bottom, CD31 immunohistochemical staining and quantification of the percentage of CD31 positive cells. Shown values are mean ± SD for n=3. Statistically significant differences between conditions are indicated with different letters. Data information: Concentrations used were 100 mg/kg/day p.o. for Palbociclib, 150 mg/kg/day p.o. for Telaglenestat, and 100 mg/kg/day p.o. Palbociclib + 150 mg/kg/day p.o. Telaglenestat for the combination. Shown values are mean ± SD for n=12 (except where stated otherwise). Significance was determined by ANOVA and Tukey’s multiple comparisons test. Statistically significant differences between conditions are indicated as *P* < 0.05 (*), *P* < 0.01 (**), and *P* < 0.001 (***).

Nevertheless, at the end of the dosage period, tumors from mice treated with Telaglenestat exhibited a reduction in their growth rate (Figure 2D-E) and about 20% of tumor growth inhibition (Figure 2G), indicating a delayed effect of this treatment. Consistent with our *in vitro* studies, mice treated with the Palbociclib and Telaglenestat combination presented the smallest tumors (Figure 2A-C) and the slowest tumor growth rates (Figure 2D-E). These results were also confirmed by bioluminescence imaging and quantification of photon radiance (Figure 2G and Supplementary Figure 2F). In addition, we observed increasing values of tumor growth inhibition with time when Palbociclib was co-administered with Telaglenestat (Figure 2F). Likewise, tumors from mice treated with the combined therapy presented the lowest proliferative activity as determined by the quantification of mitoses (mitotic count) and KI67 labeling index, and also displayed a reduction in the levels of the angiogenic marker CD31 (Figure 2H and Supplementary Figure 2G). In contrast, compared with the control group, tumors treated with Palbociclib or Telaglenestat alone did not present any changes in the levels of these markers. Strong positive correlations between the percentage of KI67 and CD31 positive cells (Pearson r = 0.828, p < 0.001), as well as between mitotic index and KI67 (Pearson r = 0.726, p < 0.01) or CD31 (Pearson r = 0.607, p < 0.04) positive cells were identified (Supplementary Figures 2H-J). Collectively, our results reveal that the Palbociclib and Telaglenestat combined treatment can effectively forestall acquired resistance to Palbociclib and significantly improve the efficacy of either treatment alone *in vivo*.

### Metabolic features of residual tumors from *in vivo* Palbociclib and Telaglenestat treatments

Next, to determine whether the tumor cells that survived the chemotherapeutic pressures have acquired sustained features of drug resistance, we collected the residual xenograft tumors at the end of the treatments for RNA sequencing (RNA-seq) and targeted metabolite profiling analyses. A total of 268 genes were differentially expressed in response to Palbociclib treatment (184 downregulated, 84 upregulated), 6,028 genes to Telaglenestat (3,290 downregulated, 2,738 upregulated), and 4,774 genes to the combined treatment (2,527 downregulated, 2,247 upregulated), compared to the control group (Supplementary Figure 3A and Supplementary Table 1). Focusing on metabolic genes, 28, 580, and 500 genes were differentially expressed between the control group and tumors treated with Palbociclib, Telaglenestat, or their combination, respectively (Supplementary Figure 3B and Supplementary Table 1). In fact, metabolism was the most affected biological process after treatment with Telaglenestat alone or in combination with Palbociclib (Supplementary Figure 3C). Tumors subjected to the dual treatment presented 581 and 87 differentially expressed metabolic genes compared to Palbociclib and Telaglenestat single treatments, respectively, implying higher metabolic differences with Palbociclib than with Telaglenestat treatment (Supplementary Figure 3D and Supplementary Table 1).

Gene Set Enrichment Analysis (GSEA) (Subramanian et al., 2005) revealed a systematic decrease in the expression levels of genes associated with metabolism, cell cycle, *MYC* oncogene, and DNA repair in tumors treated with the combination (Figure 3A), and a significant enrichment for signatures associated with protein secretion and export, suggestive of cellular senescence (Hernandez-Segura et al., 2018). Over-representation analysis (ORA) confirmed these results, showing a global reduction in the enrichment score for metabolic pathways, cell cycle, and base excision repair (Figure 3B and Supplementary Figure 4A) and a positive enrichment for senescence-related processes (Figure 3C and Supplementary Figure 4B). Consistently, the differentially expressed genes encoding enzymes of the central carbon metabolic pathways such as glycolysis, pentose phosphate pathway (PPP), tricarboxylic acid (TCA) cycle, oxidative phosphorylation (OXPHOS) and fatty acid (FA) metabolism were downregulated (Figure 3D and Supplementary Figure 5A-E). Further, the expression changes in senescence-associated genes (Supplementary Figure 5F) were in correspondence with the CellAge cellular senescence gene expression signature database (Avelar et al., 2020), indicating that the combined treatment triggers this anti-tumor mechanism leading to the irreversible arrest of cell division.

**Figure 3.**
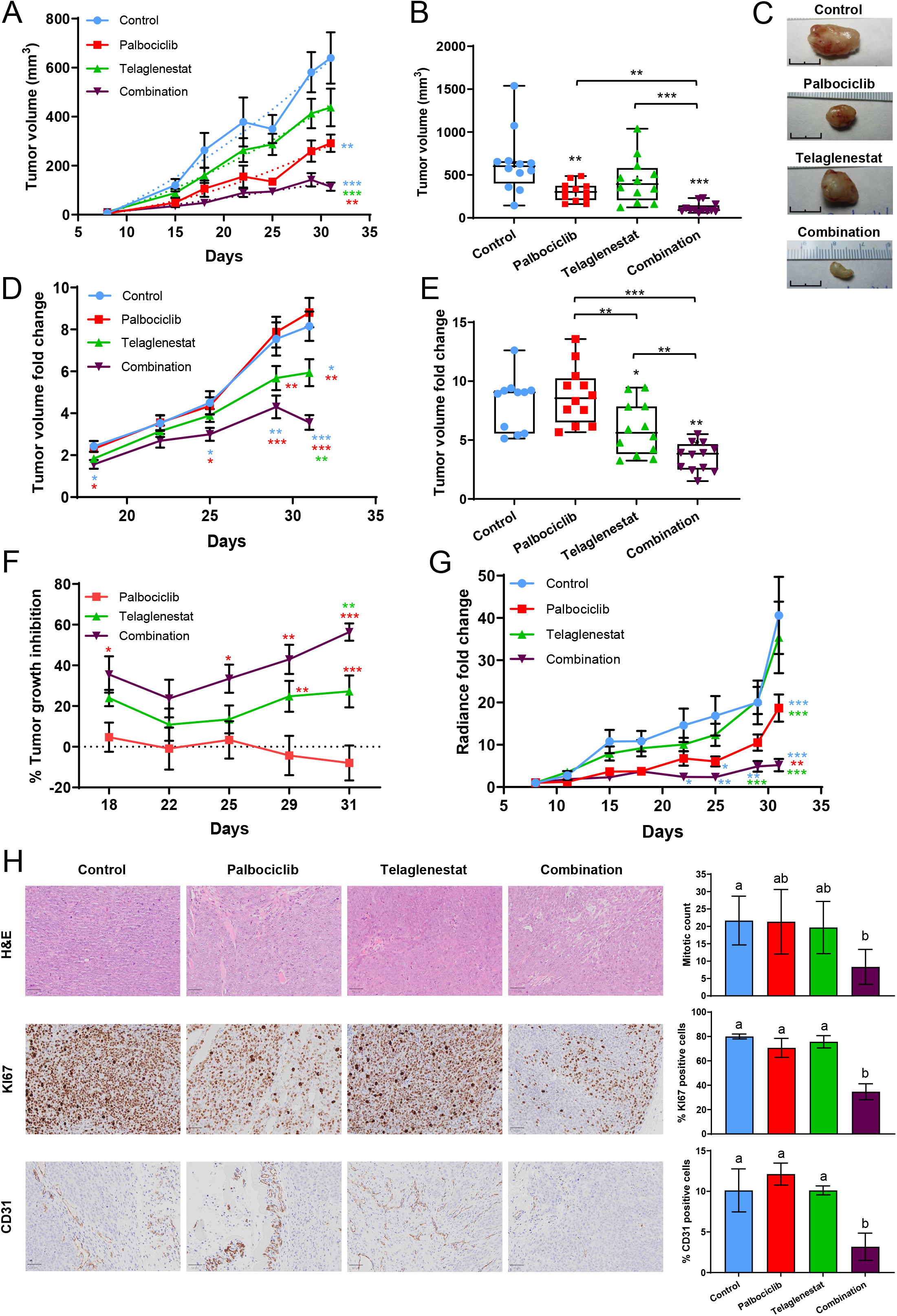
Transcriptomic analysis of Palbociclib, Telaglenestat, and their combination treatments *in vivo*. A. Gene set enrichment analysis (GSEA) results of Palbociclib and Telaglenestat *in vivo* combined treatment compared to control. RNA-Seq was performed on tumors collected after resection and immediately frozen in liquid nitrogen or isopentane. The false discovery rates *q* values (FDR) are indicated for each gene set. A positive normalized enrichment score (NES) value (bars in red) indicates enrichment in the combination treatment phenotype, while a negative NES (bars in green) indicates enrichment in the control phenotype. B, C. Over-representation analysis (ORA) of downregulated (B) or upregulated (C) gene sets in tumors treated with the combination of Palbociclib and Telaglenestat for 23 days (FDR < 0.25). Genes with FDR < 0.05 were used for the analysis. D. Schematic representation of the changes in gene expression of the central carbon metabolism caused by Palbociclib (left position in the gene label), Telaglenestat (central position), and their combination (right position) *in vivo* treatments. E, F. GSEA results of Palbociclib (E) or Telaglenestat (F) *in vivo* treatments compared to control. Gene sets significantly enriched (FDR < 0.25) are ordered by NES. A positive NES value indicates enrichment in the treatment phenotype. G, H. ORA of downregulated (G) or upregulated (H) gene sets in tumors treated with Telaglenestat for 23 days (FDR < 0.25). Data information: FDR, False discovery rate; NES, Normalized enrichment score.

In contrast, tumors treated with Palbociclib alone displayed a positive enrichment of gene signatures related to drug metabolic mechanisms, knockdown of the PTEN tumor suppressor, activation of EGFR and KRAS signaling, epithelial-mesenchymal transition, angiogenesis, and hypoxia, and a negative correlation with gene sets associated with the cell cycle, MYC targets, pyrimidine, TCA cycle and fatty acid metabolism, and DNA replication and repair (Figure 3E and Supplementary Figures 4C-D and 5G). Together, these results describe an adaptation to Palbociclib treatment towards a more aggressive tumor phenotype that differs from the adaptive response previously reported after short *in vitro* treatments (Tarrado-Castellarnau et al., 2017).

Finally, tumors from mice treated with Telaglenestat alone were negatively enriched for metabolic gene sets and exhibited an enrichment in protein secretion hallmark genes (Figure 3F-G and Supplementary Figure 4E), similarly to the dual treatment condition, while presenting increased enrichment scores for gene sets involved in the cell cycle, DNA replication and repair, and oncogenic pathways such as RAS, EGF/EGFR and PI3K (Figure 3H and Supplementary Figure 4F). These results may explain the sustained proliferation observed in tumors treated with Telaglenestat alone, as, despite a general reduction in metabolism, they acquire additional oncogenic characteristics that maintain tumor growth.

Next, we assessed the intracellular concentrations of amino acids, biogenic amines, acylcarnitines, glycerophospholipids, and sphingolipids, through targeted metabolite profiling. When compared to control, tumors subjected to the combined treatment exhibited lower concentrations of amino acids such as alanine, arginine, asparagine, aspartate, citrulline, glutamate, glycine, isoleucine, leucine, phenylalanine, proline, tyrosine, and valine, and an accumulation of glutamine resulting from the inhibition of glutaminase (Figure 4A). When comparing the four conditions, the lowest concentrations of amino acids, biogenic amines, and acylcarnitines were observed in the tumors subjected to dual treatment (Figure 4B and Supplementary Figure 6A), in consonance with the reduced metabolism inferred from the transcriptomic analysis. Principal component analysis confirmed that the drug combination caused the greatest aminoacidic (Supplementary Figure 6B) and lipidomic (Supplementary Figure 6C) shifts.

**Figure 4.**
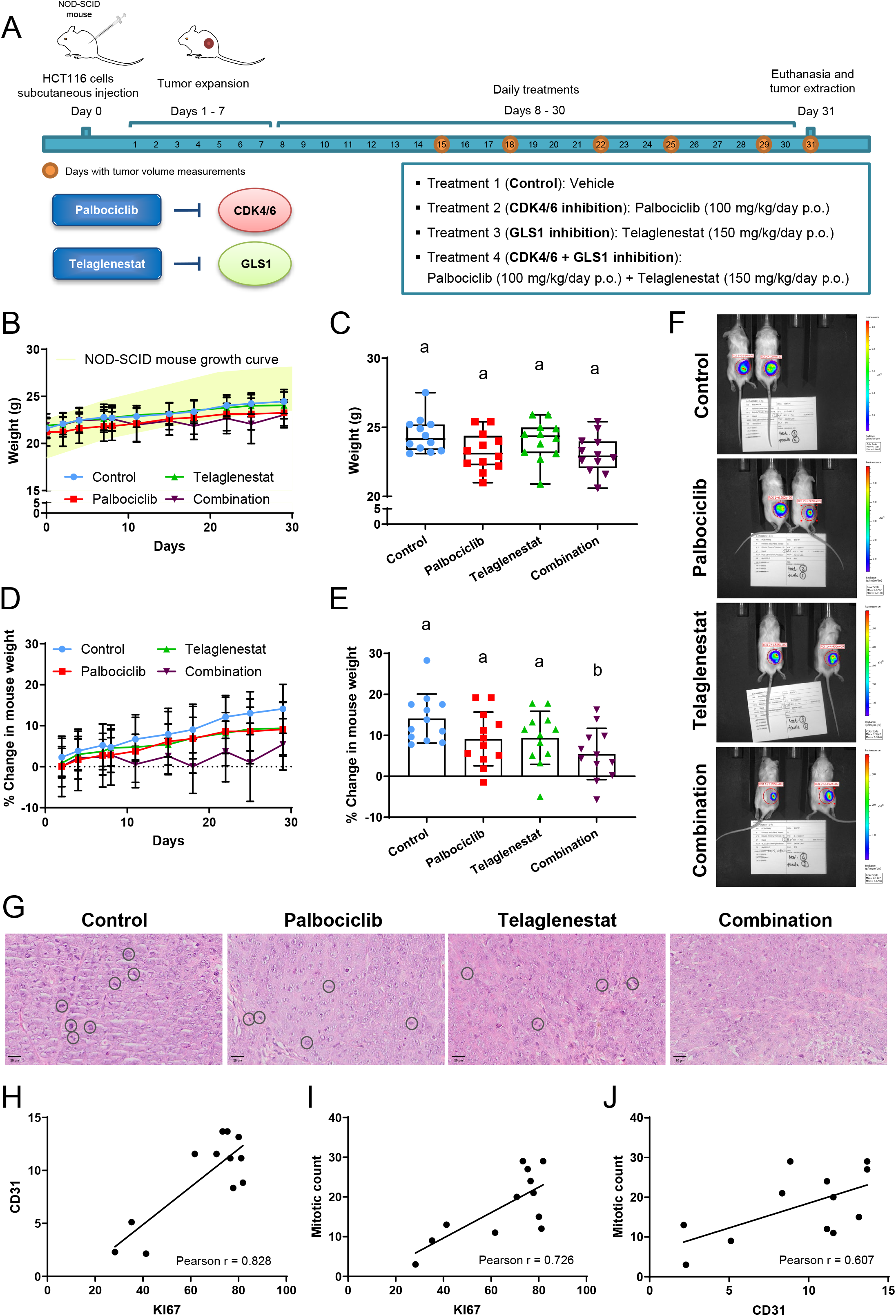
Metabolic characterization of *in vivo* tumors. A. Quantification of amino acids in tumors treated with vehicle or the combination of Palbociclib and Telaglenestat using tandem mass spectrometry coupled to liquid chromatography (LC/MS/MS). Significance was determined by a two-tailed independent sample Student’s t-tests. Statistically significant differences between treated and control cells are indicated as *P* < 0.05 (*), *P* < 0.01 (**), and *P* < 0.001 (***). B. Heatmaps of metabolite quantification profiles from tumors treated with vehicle, Palbociclib, Telaglenestat, or the combination of Palbociclib and Telaglenestat obtained by tandem mass spectrometry coupled to liquid chromatography (LC/MS/MS) or flow injection analysis (FIA/MS/MS). For phosphatidylcholines (PC), the total number of carbon atoms and double bonds of the diacyl (aa) or acyl–alkyl (ae) groups is represented by Cx:y, where x indicates the number of carbons and y the number of double bonds. The same notation is used for describing the length and the number of double bonds in the acyl chain of acylcarnitines (C), lysophosphatidylcholines (lysoPC), sphingomyelins (SM) and hydroxylated sphingomyelins (SM (OH)). C. Estimation of nitric oxide synthase (NOS), ornithine transcarbamylase (OTC), arginase (ARG), ornithine decarboxylase (ODC), spermidine synthase (SRM), and spermine synthase (SMS) enzyme activities relative to the control condition. Significance was determined by ANOVA and Tukey’s multiple comparisons test with α=0.05. Statistically significant differences between conditions are indicated with different letters. D. Schematic representation of the changes in metabolites, enzyme activities (bold), and gene expression (italics) of the polyamine metabolism and urea cycle caused by Palbociclib and Telaglenestat combined treatment. E, F. Gene set enrichment analysis (GSEA) results of Palbociclib and Telaglenestat *in vivo* combined treatment compared to palbociclib (E) and Telaglenestat (F) individual treatments. RNA-Seq was performed on tumors collected after resection and immediately frozen in liquid nitrogen or isopentane. The normalized enrichment scores (NES) are indicated for each gene set (FDR *q* value < 0.05). A positive NES value (bars in red) indicates enrichment in the combination treatment phenotype, while a negative NES (bars in green) indicates enrichment in the individual treatment condition. G. Heatmap of the expression of genes associated with Palbociclib resistance, as identified in the PALOMA-2/3 clinical trials, in tumors treated with Palbociclib, Telaglenestat, or the combination of Palbociclib and Telaglenestat, depicting the differentially expressed genes in tumors treated with the combination compared to Control group. Data information: Ala, alanine; Arg, arginine; Asn, asparagine; Asp, aspartate; Cit, citrulline; Gln, glutamine; Glu, glutamate; Gly, glycine; His, histidine; Ile, isoleucine; Leu, leucine; Lys, lysine; Met, methionine; Orn, ornithine; Phe, phenylalanine; Pro, proline; Ser, serine; Thr, threonine; Trp, tryptophan; Tyr, tyrosine; Val, valine; alpha-AAA, α-aminoadipic acid; Ac-Orn, acetylornithine; Met-SO, methionine sulfoxide; ADMA, asymmetric dimethylarginine; DMA, dimethylarginine; t4-OH-Pro, trans-4-hydroxyproline; SAM, S-adenosylmethionine; AMD, S-adenosylmethionine decarboxylase; dcSAM, decarboxylated SAM; PAOX, polyamine oxidase; SAT1, spermidine/spermine *N*^1^-acetyltransferase; ASS, argininosuccinate synthase; ASL, argininosuccinate lyase. Shown values are mean ± SD for n=4.

Conversely, we found the highest levels of phosphatidylcholines and sphingomyelins in tumors treated with the Palbociclib and Telaglenestat combination (Figure 4B and Supplementary Figure 6D-E), suggesting that cells accumulate lipids as essential components of membranes but are unable to duplicate due to the alterations of DNA synthesis, resulting in membrane remodeling. Further, in tumors treated with the drug combination, estimation of enzyme activities through metabolite ratios (Dossus et al., 2021) revealed a decrease in the activity of stearoyl CoA-desaturase (SCD) (Supplementary Figure 6F), involved in cell migration and invasion (Ran et al., 2018), and in the urea cycle and spermine synthase (SMS) of the polyamine pathway (Figure 4C-D and Supplementary Figure 6G), in line with the observed downregulation of MYC target genes (Figure 3A) and the reduced expression of genes involved in tumor invasion and metastasis (Gerner and Meyskens, 2004). The accumulation of spermidine (Supplementary Figure 6G) is associated with autophagy induction, which suppresses proliferation and promotes apoptosis in cervical cancer cells (Chen et al., 2018), and could explain the enrichment observed in autophagic processes in tumors treated with the combination (Figure 3C).

The combined treatment also elicited a downregulation of arginine methylation (Supplementary Figure 6H) and protein arginine methyltransferases (PRMTs) (Supplementary Figure 6I), which orchestrate epigenetic modifications, chromatin architecture, transcription, signaling, splicing, DNA damage response, and cell metabolism, and are identified as promising cancer therapeutic targets (Wu et al., 2021). In fact, evaluation of the transcriptomic data from the PALOMA-2 (NCT01740427) and PALOMA-3 (NCT01942135) clinical trials for Palbociclib (Zhu et al., 2022) evidences a significant positive correlation between the expression of both *PRMT1* and *CARM1* genes and the overexpression of genes in glycolysis, OXPHOS, hypoxia, PI3K/AKT/mTOR and MYC signaling pathways (Supplementary Figure 6J and Supplementary Table 2), which are associated with the acquisition of therapeutic resistance and shorter progression-free survival in the PALOMA-2/3 trials. Likewise, *PRMT1* and *CARM1* are positively correlated with the expression of Palbociclib resistance genes, while they are negatively correlated with the sensitivity genes identified in the PALOMA-2/3 trials comparative biomarker analyses (Zañudo et al., 2022; Zhu et al., 2022) (Supplementary Table 3). These observations suggest that residual tumor cells that survive Palbociclib therapy have acquired chemoresistance features through an epigenetic mechanism. Indeed, the combination of Palbociclib with the selective inhibitor of type I PRMTs GSK3368715 (Fedoriw et al., 2019) caused a synergistic reduction of cell proliferation in HCT116, SW403, HT29, and SW620 colorectal cancer cell lines over a wide dose range (Supplementary Figure 6K-N), regardless of their CMS subtype or microsatellite stability status. Furthermore, Palbociclib long-term resistant HCT116 cells, which displayed 4-fold higher Palbociclib IC_50_ values compared to control HCT116 cells (Supplementary Figure 6O), exhibited increased sensitivity to the inhibition of type I PRMTs, with 5-fold lower IC_50_ values (Supplementary Figure 6P), indicating that Palbociclib resistance may also be overcome by direct type I PRMT inhibition and providing a potential explanation for the reported efficacy of the combined treatment of Palbociclib and Telaglenestat.

On the other hand, tumors subjected to the combined treatment displayed a downregulated expression of metastasis, cell cycle progression, and MYC and mTOR oncogenic pathways, as well as an increase in protein secretion and senescence pathways compared to the single treatments (Figure 4E-F). Consistently, in tumors treated with the combination, the expression of Palbociclib resistance genes identified in PALOMA-2/3 (Zañudo et al., 2022; Zhu et al., 2022) and PEARL (NCT02028507) (Guerrero-Zotano et al., 2023; Martin et al., 2021) trials was significantly decreased (Figure 4G and Supplementary Figure 6Q). Indeed, Palbociclib-resistant genes identified in the PALOMA-2/3 and PEARL trials were enriched in the genes significantly downregulated after combined treatment, with Fisher exact test P-values of 0.00113 (Odds ratio = 3.06) and 0.00283 (Odds ratio = 2.98), respectively.

### Genome-scale metabolic modeling reveals that Telaglenestat is uniquely suited to counter the adaptive metabolic reprogramming induced by Palbociclib

To obtain a system-wide perspective of the metabolic reprogramming underlying treatment with Palbociclib, Telaglenestat, or their combination, we integrated the transcriptomics and metabolomics data from the residual xenograft tumors in the framework of Recon3D. Recon3D is a genome-scale reconstruction of human metabolism, providing a mathematical representation of all the reactions and transport processes known in human cells and the enzymes and transmembrane carriers mediating them (Brunk et al., 2018). More in detail, we used a quadratic multiomic metabolic transformation algorithm (qM^2^TA) to impute the metabolic transformation underlying exposure to each drug and their combination by finding the intracellular metabolic flux rewiring most consistent with the transcriptomics and metabolomics measures (Figure 5A and Supplementary Figure 7A). Treatment with Palbociclib resulted in 215 reactions with a flux value increase of 25% or more relative to control, whereas 239 had their flux reduced by 25% or more. Conversely, Telaglenestat and the combined treatment resulted in a larger number of reactions with reduced flux values relative to the control, 1,461 and 1,211, respectively, whereas only 726 and 701 reactions increased their flux values by 25% or more in such treatments (Supplementary Table 4). This suggests that Telaglenestat, alone or in combination, tends to have a largely inhibitory effect on metabolism, a notion also apparent by observing the distribution of reaction flux values relative to the control (Figure 5B).

**Figure 5.**
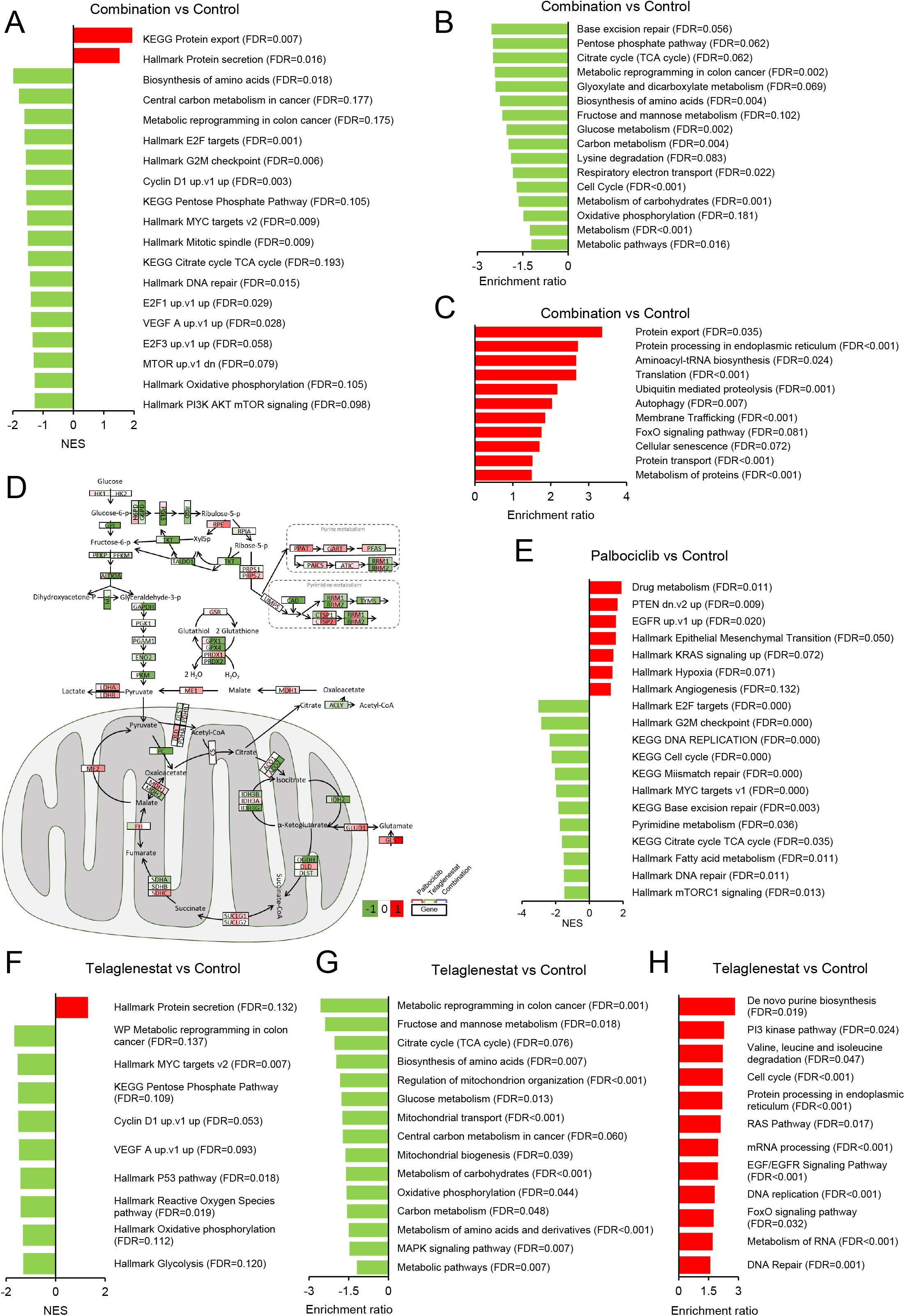
Quadratic Multiomic Metabolic Transformation Algorithm (qM^2^TA). A. Schematic representation of qM^2^TA. qM^2^TA simulates the metabolic flux transition from the control condition to treatment with Palbociclib, Telaglenestat, or Palbociclib and Telaglenestat through the integration of transcriptomics and metabolomics into a genome-scale metabolic model (GSMM). A hypothetic 3D plot of three flux values (v_x_,v_y_,v_z_) and their transition with each treatment is provided. B. Box plot of log2 fold changes in flux reaction values with each treatment relative to control. Dotted red lines indicate fold changes over 1.25 and under 0.75. C. Variations in pathways of central carbon and oxidative metabolism with each treatment. Results are expressed as log2 fold changes for total pathway flux relative to the control.

Analyzing the flux through pathways of central carbon metabolism and OXPHOS, both key for cellular proliferation and the most relevant by the magnitude of carried flux, offers a more nuanced picture (Figure 5C and Supplementary Table 4). For instance, Palbociclib treatment caused an increased flux through oxidative phosphorylation reactions and a decreased flux through purine and pyrimidine synthesis reactions. Indeed, activation of OXPHOS can likely be attributed to increased mTOR and MYC signaling, given that both promote mitochondrial biogenesis (Cunningham et al., 2007; Morita et al., 2013; Morrish and Hockenbery, 2014). Conversely, Telaglenestat reduced the flux through the OXPHOS, glycolysis, and pentose phosphate pathways while strongly upregulating purine and, to a lesser extent, pyrimidine metabolism. Finally, the combination of Palbociclib and Telaglenestat offered the best of each individual treatment with no significant upregulation of OXPHOS and a reduced flux through the pathways of purine and pyrimidine synthesis, glycolysis, and pentose phosphate pathway.

Upregulated pathways after treatment with Palbociclib or Telaglenestat likely reflect a process of metabolic reprogramming that allows cancer cells to adapt to and counter drug-induced stress. With this in mind, we used qM^2^TA to systematically search for potentially targetable enzymes or transmembrane carriers to prevent the metabolic adaptation to either Palbociclib or Telaglenestat (Supplementary Figure 7B). We identified 327 and 280 potential targets to partially revert the metabolic adaptation underlying the Palbociclib or Telaglenestat treatments, respectively (Supplementary Table 4). It is worth mentioning that glutaminase, the therapeutic target of Telaglenestat, was one of the targets against Palbociclib adaptation identified in this analysis, in accordance with our previous studies (Tarrado-Castellarnau et al., 2017). Even more remarkable, 112 targets inferred against Palbociclib adaptation, including many top-scoring targets, corresponded to genes that were significantly downregulated in response to Telaglenestat. Likewise, Palbociclib treatment downregulated 8 of the putative targets identified against Telaglenestat adaptation, including the second top-scoring putative target, ribonucleotide reductase. Given that only 15 metabolic genes were significantly downregulated by Palbociclib, putative targets against Telaglenestat-induced metabolic adaptation were significantly overrepresented (Fisher exact test P-value = 0.01319 and Odds ratio = 3.83) (Supplementary Table 5). These findings are in line with the patterns that emerged upon observation of flux changes through central carbon metabolism pathways (Figure 5C). Indeed, they reinforce the notion that Palbociclib and Telaglenestat are uniquely suited to counter the metabolic reprogramming induced by the reciprocal drug, thus providing a mechanistic rationale for the high effectiveness of the drug combination observed *in vitro* and *in vivo*.

### Cells from residual tumors treated *in vivo* with Palbociclib and Telaglenestat maintain persistent drug-specific metabolic reprogramming *ex vivo*

Given that the effects of Palbociclib and Telaglenestat are time-dependent, we determined the potential persistence of their adaptive or resistant phenotypes after *in vivo* chemotherapy. To this end, we explanted 12 fresh residual tumors from mice treated for 23 days with vehicle, Palbociclib, Telaglenestat, or their combination, obtaining a total of 12 distinct two-dimensional monolayer cell cultures (three per *in vivo* treatment condition). Once established, we analyzed the glycolytic and glutaminolytic profiles of each cell line by incubation with complete media without any chemotherapeutic pressure. We found that cells derived from tumors that had been treated with the drug combination exhibited a decrease in the metabolic fluxes of glucose, glutamine, lactate, and glutamate (Figure 6A), together with a reduction in proliferation and doubling time (Figure 6B and Supplementary Figure 8A) without affecting cell cycle distribution (Supplementary Figure 8B). Likewise, the overall consumption rate of hexoses was only significantly reduced in the condition resulting from tumors treated with Palbociclib and Telaglenestat (Figure 6C). In addition, we assessed the amino acid exchange fluxes and found that alanine production, and arginine, asparagine, aspartate, histidine, isoleucine, leucine, lysine, methionine, phenylalanine, proline, and valine consumption rates were diminished in cells from tumors that had been dosed with the combination therapy (Figure 6D and Supplementary Figure 8C). To further confirm the lower glycolytic rates observed, we measured in real-time the extracellular acidification rate (ECAR), essentially a sign of lactate production from glycolysis with a small contribution from the CO_2_ produced in other reactions, while sequentially adding glucose and oligomycin to cells cultured in a medium without glucose. As expected, cells derived from Palbociclib plus Telaglenestat-treated tumors portrayed a significant reduction in the glycolytic capacity, glycolytic reserve, and non-glycolytic acidification (Figure 6E). Then, we confirmed that the metabolic downregulation caused by the long-term double treatment and reported *in vivo* and 2D *ex vivo* cultures was also maintained in spheroid cultures (Figure 6F).

**Figure 6.**
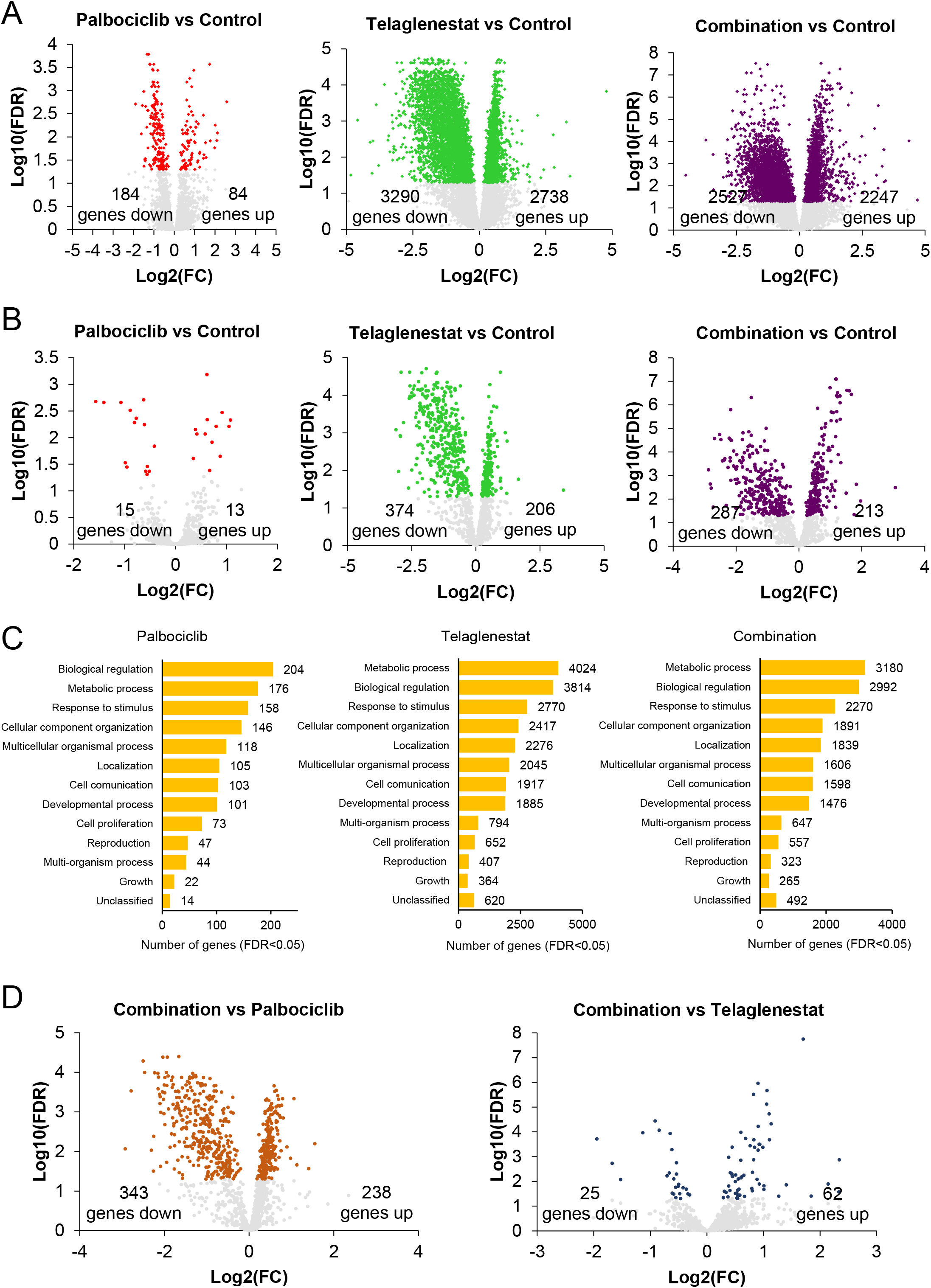
Metabolic characterization of cell lines derived from HCT116 xenografts. Twelve cell lines were obtained from HCT116 tumors from mice that had been treated daily with vehicle, Palbociclib, Telaglenestat, or the combination of Palbociclib and Telaglenestat for 23 days (three cell lines per condition). All cell lines were grown in the absence of chemotherapeutics. A. Comparative extracellular metabolic fluxes. Glucose and glutamine consumption and lactate and glutamate production rates were assessed after 24 h of incubation with fresh media and normalized to cell number. B. Cell proliferation curves. Results are shown as fold change of cell proliferation relative to the measurement at 24 h (mean ± SD of n=6). Significance was determined by one-way ANOVA and Tukey’s multiple comparisons test. Statistically significant differences between cells derived from tumors that had been treated with the combination and cells obtained from control tumors are indicated as *P* < 0.05 (*). C. Pool of hexoses consumption rate determined by targeted metabolomics after 24 h of incubation with fresh media and normalized to cell number. D. Amino acid consumption and production rates were measured after 24 h of incubation with fresh media and normalized to cell number. Significance was determined by a two-tailed independent sample Student’s t-test. Statistically significant differences between treated and control cells are indicated as *P* < 0.05 (*), *P* < 0.01 (**), and *P* < 0.001 (***). E. Quantification of extracellular acidification rate (ECAR) for glycolysis, glycolytic capacity, glycolytic reserve, and non-glycolytic acidification. Cells were cultured in the absence of glucose, and sequential injections of glucose and oligomycin were applied. Data are normalized to cell number and represented as the mean of three independent cell lines for each condition ± SD with n=5. F. Spheroids comparative extracellular metabolic fluxes. Spheroids were grown in non-adherend conditions for each cell line, and glucose and glutamine consumption and lactate and glutamate production rates were assessed after 96 h of incubation with fresh media and normalized to cell number. G. Quantification of oxygen consumption rates (OCR), following sequential injections of oligomycin (1 µM), CCCP (600 nM), and antimycin A (2 µM) and rotenone (2 µM), for non-mitochondrial oxygen consumption, basal respiration, maximal respiration, non-ATP linked oxygen consumption (proton leak), ATP production-associated respiration, and spare respiratory capacity. Data are normalized to cell number and shown as the mean of three independent cell lines for each condition ± SD with n=5. H. Western blotting analysis of total protein fractions for the cell lines generated from tumors using β-actin as a loading control. Each lane corresponds to a different cell line. I. Quantification of the ATP production rate from glycolysis (glycoATP) and mitochondrial oxidative phosphorylation (mitoATP) and their contribution to the total cellular ATP production rate. Data are normalized to cell number and displayed as the mean of three independent cell lines for each condition ± SD with n=5. J. Energetic map exhibiting the distribution of mitochondrial ATP (mitoATP) production rate vs. glycolytic ATP (glycoATP) production rate. Cells derived from tumors treated with the combination of Palbociclib and Telaglenestat presented the most quiescent metabolism. Values are expressed as the mean of three independent cell lines for each condition ± SD with n=5. K. IC_50_ values for palbociclib and the combination of Palbociclib and Telaglenestat in cell lines derived from tumors that had been treated with these inhibitors *in vivo*. Bars represent the mean of the IC_50_ values of three independent cell lines for each condition ± SD with n=6. Data information: Ala, alanine; Arg, arginine; Asn, asparagine; Asp, aspartate; Cit, citrulline; Gln, glutamine; Glu, glutamate; Gly, glycine; His, histidine; Ile, isoleucine; Leu, leucine; Lys, lysine; Met, methionine; Orn, ornithine; Phe, phenylalanine; Pro, proline; Ser, serine; Thr, threonine; Trp, tryptophan; Tyr, tyrosine; Val, valine. Significance was determined by one-way ANOVA and Tukey’s multiple comparisons test with α=0.05. Statistically significant differences between conditions are represented with different letters. Data are shown as the mean of three independent cell lines for each condition ± SD with n=3 (except otherwise indicated).

Considering that the TCA cycle and OXPHOS were among the gene sets downregulated *in vivo* by the treatments (Figure 3), we performed a pharmacological profiling of the mitochondrial respiratory function by combining the ATP synthase inhibitor oligomycin, the protonophoric uncoupler CCCP, the complex III inhibitor antimycin A and the complex I inhibitor rotenone, while measuring the oxygen consumption rate (OCR). This analysis revealed that cells explanted from tumors treated with the drug combination presented lower non-mitochondrial oxygen consumption, basal respiration, maximal respiration, non-ATP linked oxygen consumption (proton leak), ATP production-associated respiration, and spare respiratory capacity than control cells, depicting a global reduction in mitochondrial respiratory capacity (Figure 6G), consistent with the reported lower consumption rates of glucose and amino acids and the decrease of MYC, p-AKT, KGA and GLDH protein levels (Figure 6H). Moreover, RNA-seq analysis revealed that OXPHOS, one of the main pathways enriched during metastatic seeding and chemoresistance (Davis et al., 2020; El-Botty et al., 2023; Sica et al., 2020), was downregulated in cells derived from the combination therapy tumors compared to those obtained from control or single-agent treated tumors (Supplementary Figure 8D).

### Combined Palbociclib and Telaglenestat treatment induces a less aggressive phenotype without the acquisition of resistance

Next, to compare the cellular function and energy demands of the surviving cancer cells after each chemotherapeutic treatment, we measured the ATP production rate *ex vivo* as a more accurate approach to assess the energetic cellular state than determining the total intracellular levels of ATP, which are maintained under steady-state conditions (Hardie et al., 2012). We observed that cells from tumors that were treated with the combined therapy displayed the lowest rate of mitochondrial, glycolytic, and total ATP production, while the reduction in both single-agent treatments was less pronounced (Figure 6I). Similarly, the contribution of mitochondria to oxygen consumption and proton production was significantly lower in cells previously treated *in vivo* with the combination (Supplementary Figure 8E-F), consistent with a quiescent phenotype (Figure 6J). Of note, xenograft-explanted cells presented a shift from glycolytic to mitochondrial metabolism compared to *in vitro* HCT116 cells not passaged in mice, which had an ATP rate index of 1 (Figure 6I and Supplementary Figure 8G). These results agree with the observed reduction in intracellular glutathione in cells derived from tumors treated with the combination or Telaglenestat alone (Supplementary Figure 8H), considering that this antioxidant molecule is synthesized through two ATP-requiring steps from glutamate.

To complement this analysis, we performed GSEA, which showed that cells explanted from tumors subjected to the combined therapy exhibited a downregulation of genes involved in the cell cycle machinery, MYC signaling, metastasis, OXPHOS, and electron transport chain, as compared to cells derived from tumors treated with Palbociclib alone (Supplementary Figure 8I). These cells also exhibited a downregulation of genes regulated by oncogenes such as MYC, VEGF, and cyclin D1 and associated with mitochondria and amino acid metabolism when compared with cells from tumors treated with Telaglenestat alone (Supplementary Figure 8J), revealing a persistent reduction in the malignant potential of surviving cells that had been exposed to combined therapy *in vivo*, as compared to those subjected to a single agent.

Finally, to examine whether the xenograft-derived cell lines had shifted their sensitivity profiles after *in vivo* exposure to the therapies, we exposed the cells to increasing doses of each drug or the combination for 96 h and determined IC_50_ values. While the effect of Telaglenestat on cell proliferation was similar in all cases, regardless of *in vivo* drug exposure (Supplementary Figure 8K), *in vivo* exposure to Palbociclib alone or in combination with Telaglenestat conferred explanted tumor cells with a 2-fold increase in resistance to Palbociclib alone (Figure 6K). Remarkably, the same cells were at least as sensitive as control HCT116 cells to the combination of Palbociclib and Telaglenestat (Figure 6K). Moreover, the combination caused *in vitro* synergistic antiproliferative effects in the cell lines derived from tumors subjected to the combined therapy *in vivo* (Supplementary Table 6).

Together, these data demonstrate that long-term Palbociclib and Telaglenestat combined treatment impairs glycolytic metabolism, mitochondrial respiration, and ATP production, as well as mitigates oncogenic pathways, in colon cancer cells that survive *in vivo* therapy with both drugs, resulting in a persistently less aggressive tumor phenotype.

## Discussion

Targeting cell cycle through cyclin-dependent kinases 4 and 6 (CDK4/6) inhibition (CDK4/6i) is a promising strategy in cancer therapy since they play a major oncogenic role in a wide range of human cancers (Goel et al., 2022). However, few cancer types are sensitive to CDK4/6i’s when used as single agents (Álvarez-Fernández and Malumbres, 2020; Herrera-Abreu et al., 2016). Furthermore, sensitive cancers invariably acquire resistance to CDK4/6 inhibition (Papadimitriou et al., 2022), such that CDK4/6i’s are only efficacious in combination with drugs that target other pathways on which cancer cells rely to escape CDK4/6 dependency (Gennari et al., 2021; Ma et al., 2023). Numerous pre-clinical studies and clinical trials are currently exploring the therapeutic potential of combining CDK4/6 inhibitors with endocrine therapies (e.g., aromatase inhibitors, fulvestrant), immunotherapies (e.g., antibodies to PD1/PD-L1), oncogenic kinase inhibitors (e.g., receptor tyrosine kinase-PI3K-AKT-mTOR signaling, RAS, BRAF, MEK, IGF-1R, ALK inhibitors), and other targeted or chemotherapeutic agents (e.g., autophagy, PARP inhibitors) in solid tumors (Goel et al., 2022).

Metabolic reprogramming is a relevant cell-autonomous mechanism through which tumor cells change their energetic dependencies to alternative pathways in order to bypass the selective pressures exerted by a given class of antineoplastic drugs (Mendez-Lucas et al., 2020; Pranzini et al., 2021). This metabolic shift may reveal vulnerabilities that can be targeted through combination therapies to prevent and overcome chemoresistance (El-Botty et al., 2023; Labrie et al., 2022). Our previous work identified the inhibition of glutaminase as a metabolism-based strategy to combine with CDK4/6 targeting therapy (Tarrado-Castellarnau et al., 2017), a combination subsequently corroborated as effective to overcome CDK4/6i resistance in esophageal squamous cell carcinoma (Qie et al., 2019) and lung cancer (Conroy et al., 2020). Here, we have undertaken an in-depth, unbiased metabolomics approach to assess in greater detail the metabolic reprogramming undergone by HCT116 colorectal cancer cells exposed to either the CDK4/6i Palbociclib, the glutaminase inhibitor Telaglenestat, or their combination.

First, we observed that the metabolic reprogramming undergone by cancer cells after treatment with either agent alone is time-dependent. Consistently, tumors in mice dosed with Palbociclib alone had the same growth rate as the control tumors despite exhibiting a significant reduction in their volume, suggesting that Palbociclib activity is also time-dependent *in vivo* and that, after the initial antiproliferative effect, surviving tumor cells acquire the capacity to overcome this inhibition and restore their proliferative rate. Telaglenestat did not exert any significant effect on tumor growth in xenotransplanted mice. In contrast, the synergistic antiproliferative effect of the dual treatment was time-independent and effective both *in vitro* and *in vivo*. As such, the combined administration of Palbociclib and Telaglenestat was strongly synergistic for the inhibition of tumor growth, proliferative index, and angiogenesis.

Gene set enrichment analysis confirmed that tumors subjected to the double therapy exhibited a downregulation in metastasis, cell cycle progression, and MYC and mTOR oncogenic pathways compared to control and single treatments. Lending translational significance to these observations, tumors from mice subjected to dual therapy presented a decrease in the expression of Palbociclib-resistance genes identified in the PALOMA-2/3 and PEARL clinical trials assessing biomarkers of sensitivity and resistance to palbociclib in ER^+^ HER^-^ breast cancer (Guerrero-Zotano et al., 2023; Martin et al., 2021; Zañudo et al., 2022; Zhu et al., 2022). Of particular interest, the treatment of HCT116 tumor-bearing mice with Palbociclib and Telagleneastat provoked the downregulation of arginine methylation and protein arginine methyltransferases (PRMTs). Expression of PRMTs correlates with the activation of resistance pathways associated with lower progression-free survival (Zhu et al., 2022) and is upregulated through MYC and mTORC1 activation (Litzler et al., 2023), suggesting that Palbociclib-surviving cells might have acquired chemoresistance through an epigenetic mechanism.

The drug-specific metabolic reprogramming of residual cancer cells after *in vivo* treatments was maintained in *ex vivo* cultures for many cell generations, in the absence of any chemotherapeutic selective pressure, also suggestive of a heritable epigenetic mechanism underlying the persistent metabolic reprogramming. Specifically, cells derived from tumors treated with the Palbociclib and Telaglenestat combination presented a general decrease in central carbon metabolic fluxes in conjunction with an overall reduction in mitochondrial respiration, as well as in mitochondrial, glycolytic, and total ATP production. Notably, these cells displayed an impoverishment in oncogenic pathways compared to cells from tumors subjected to Palbociclib or Telaglenestat alone, indicating a less aggressive phenotype. Importantly, residual tumor cells from *in vivo* dual treatment were sensitive to the double treatment *ex vivo*. In contrast, residual tumor cells from *in vivo* treatment with Palbociclib as a single agent were resistant to this drug *ex vivo*. This indicates that (i) Palbociclib treatment readily elicits a resistant phenotype both *in vitro* and *in vivo*, (ii) the combination of Palbociclib with Telaglenestat disables Palbociclib-induced resistance mechanisms *in vitro* and *in vivo* and (iii) the disabling of resistance mechanisms by this drug combination is heritable through many cell generations.

By applying the quadratic multiomic metabolic transformation algorithm (qM^2^TA), we integrated metabolite and enzyme transcript levels to infer genome-scale metabolic rewiring upon treatment with Palbociclib, Telaglenestat, or their combination *in vivo*. From this analysis, it emerged that Palbociclib treatment resulted in an increased flux through OXPHOS, attributable to increased mTOR and MYC signaling (Cunningham et al., 2007; Morita et al., 2013; Morrish and Hockenbery, 2014), and decreased flux through reactions involved in nucleotide synthesis. OXPHOS is often upregulated in cancer stem cells and drug-resistant tumors (Davis et al., 2020; Sica et al., 2020) and our observations are in line with recent findings describing that the treatment of metastatic breast cancer with Palbociclib elicits increased OXPHOS as a resistance mechanism that can be countered pharmacologically (El-Botty et al., 2023). In contrast to Palbociclib, Telaglenestat treatment caused a decreased flux through OXPHOS, glycolysis, and the pentose phosphate pathway, while increasing flux through nucleotide metabolism. Notably, nucleotide synthesis, pentose phosphate pathway, glycolysis, and TCA play a key role in the synthesis of biomass building blocks needed for proliferation and are commonly upregulated in cancer compared to healthy tissue (Tarrado-Castellarnau et al., 2016). Hence, the combination of Palbociclib and Telaglenestat maintained the benefits of each individual treatment while sparing the upregulation of OXPHOS as a resistance mechanism and reducing flux through nucleotide synthesis, glycolysis, and pentose phosphate pathways. Furthermore, many markers of metabolic adaptation to Palbociclib were downregulated by Telaglenestat and *vice versa*. Taken together, our analysis indicates that the upregulation of OXPHOS through glutaminolysis is the most relevant mechanism of resistance to CDK4/6 inhibition in our cell model. As such, inhibition of glutaminolysis through Telaglenestat, or other selective GLS1 inhibitors, is an optimal strategy to overcome resistance to Palbociclib in cancer cells.

In summary, we have delineated a major metabolic mechanism explaining resistance to Palbociclib in a colorectal cancer model, namely glutaminolysis-driven OXPHOS, and shown that the combination of a selective CDK4/6 inhibitor with a selective glutaminase inhibitor optimally resensitizes cancer cells to CDK4/6 inhibition through many cell generations. We believe that these findings warrant a proposal to use this drug combination in clinical settings.

## Acknowledgments

We are grateful for assistance from the Experimental Histopathology and High Throughput Screening facilities at the Francis Crick Institute. This work was supported by the Spanish Ministry of Science and Technology (MCIU/AEI/FEDER, UE; SAF2017-89673-R, MCIN/AEI/10.13039/501100011033; PID2020-115051RB-I00 and PID2019-107139RB-C21), the Interdisciplinary Platform-Global Health (Plataforma Temática Interdisciplinar-Salud Global, PTI-SG; SGL2103019), the Networked Researched Center on Liver and Digestive Diseases (Centro de Investigación en Red en Enfermedades Hepáticas y Digestivas, CIBER-EHD; EHD20PI03 and CB17/04/00023), the Catalan Agency for Universities and Research (AGAUR) (2021-SGR-00350 and 2021-SGR-01490), ICREA Foundation (ICREA Academia award granted to M. Cascante) and The Francis Crick Institute, which receives its core funding from Cancer Research UK (FC001223), the UK Medical Research Council (FC001223) and the Wellcome Trust (FC001223).

## Author contributions

M.T-C., M.Y., T.M.T, and M.C. conceived and designed the study. M.T-C. led and performed experiments with J.T-C. M.P., C.H., J.P., E.Z., and I.H.P. assisted with experiments. C.F. and J.J.L. carried out bioinformatics analyses. M.T-C., C.F., T.M.T., and M.C. analyzed the data. A. S-B. analyzed histopathology images. S.M. set up the targeted metabolomics method. M.T-C. and C.F. wrote the manuscript. M.T-C., C.F., J.T-C., M.Y., T.M.T, and M.C. reviewed and revised the manuscript. All authors approved the final manuscript. M.T-C., M.Y., T.M.T, and M.C. secured funding and supervised the study.

## Declaration of interests

1. H. P. is a current employee and stockholder of BioNTech SE, Mainz, Germany, which had no involvement in this work financially or otherwise. The other authors declare no potential conflicts of interest.

## Methods

### Compounds and reagents

Palbociclib (PD0332991) and Telaglenestat (CB-839) were purchased from Selleckchem (Houston, TX, USA). The glutaminase inhibitor bis-2-(5-phenylacetamido-1,2,4-thiadiazol-2-yl)ethyl sulfide (BPTES) was kindly provided by Dr. Mariia Yuneva (The Francis Crick Institute, London, UK). The inhibitor of type I PRMTs GSK3368715 was purchased from Cayman Chemical (Ann Arbor, MI, USA). Stock solutions of 10 mM were prepared with water or dimethyl sulfoxide (DMSO) according to the manufacturer’s instructions. Antibiotics (10,000 U/ml penicillin, 10 mg/ml streptomycin), PBS, and Trypsin EDTA solution C (0.05% trypsin – 0.02% EDTA) were obtained from Biological Industries (Kibbutz Beit Haemet, Israel), and fetal bovine serum (FBS) from Invitrogen (Carlsbad, CA, USA).

### Cell culture

The human colorectal carcinoma cell lines HCT116, SW403, HT29, and SW620, and the human embryonic kidney 293T (HEK293T) cells were obtained through the American Type Culture Collection (ATCC, Manassas, VA, USA). The HCT116 derivative HCT116^Luciferase-GFP^ cells were obtained through lentiviral transfection and transduction. Twelve HCT116 derivative cell lines were obtained from tumors from mice that had been treated daily with vehicle, Palbociclib, Telaglenestat, or the combination of Palbociclib and Telaglenestat for 23 days (three independent cell lines per condition). All cell lines were grown in the absence of chemotherapeutics. Palbociclib long-term resistant cell line was established by culturing HCT116 cells in gradient concentrations of Palbociclib for four months. HCT116 cells and all the derivative cell lines were cultured in Dulbecco’s modified Eagle medium (Gibco, Thermo Fisher Scientific Inc., Waltham, MA, USA) / Nutrient mixture HAM F12 (Biological Industries) (DMEM/F12, 1:1 mixture) with 2.5 mM L-glutamine and 12.5 mM D-glucose. SW403, HT29, SW620, and HEK293T cells were cultured in DMEM culture medium. Media were supplemented with 10% FBS, penicillin (50 U/ml), and streptomycin (50 µg/ml). Cells were incubated at 37 °C in a humidified atmosphere with 5% CO_2_. All cell lines were regularly tested for mycoplasma contamination and cultivated for less than three months after receipt, generation, or thawing.

### Lentiviral transfection and transduction

HEK293T cells were transiently transfected using Fugene reagent (Roche, Indianapolis, IN, USA) with pCMV-GFP/Luc plasmid (kindly provided by Dr. Timothy M. Thomson) for the constitutive co-expression of the firefly luciferase gene and GFP. Viral supernatants were harvested 48 h after transfection, filtered through 0.45 μm methylcellulose filters (Merck Millipore, Billerica, MA, USA), and used to transduce HCT116 recipient cells. 5×10^5^ HCT116 cells were cultured in the presence of the viral supernatant and 8 μg/ml polybrene for 24 h. GFP-positive transduced cells were selected by fluorescence-activated cell sorting (FACS) (MoFlo, Beckman Coulter). Luminescence levels yielded by cell lysates were measured using the Luciferase Assay System kit (Promega, Madison, WI, USA) in a Mithras LB 940 plate reader (Berthold Technologies, Bad Wildbad, Germany).

### Cell proliferation assays

Having optimized the seeding density, 2×10^3^ cells per well were seeded in 96-well plates, and media were replaced at 24 h with fresh media containing different concentrations of each drug, the combination of drugs, or vehicle. Plates were placed in an IncuCyte® S3 (Sartorius, Göttingen, Germany) live-cell analysis system to monitor cell confluence. At 96 h, cells were washed with PBS and lysed with 0.01% SDS, and plates were stored frozen at -20 °C overnight. Then, plates were thawed at 37 °C and incubated in darkness with 4 µg/ml Hoechst (HO33342; 2’-[4-ethoxyphenyl]-5-[4-methyl-1-piperazinyl]-2,5’-bi-1H-benzimidazole trihydrochloride trihydrate, Sigma-Aldrich, St Louis, MO, USA) DNA stain in solution buffer (1 M NaCl, 1 mM EDTA, 10 mM Tris-HCl pH 7.4) on a shaker at 37 °C for 1 h. Finally, fluorescence was measured in a FLUOstar OPTIMA microplate reader (BMG LABTECH GmbH, Ortenberg, Germany) at 355 nm excitation and 460 nm emission. Cell proliferation was represented as a percentage relative to untreated control cells. Concentrations that caused 50% of inhibition of cell proliferation (IC_50_) were calculated using GraphPad Prism 7 software (GraphPad Software, San Diego, CA, USA). Cell size and volume values were obtained using a Scepter^TM^ Handheld Automated Cell Counter (EMD Millipore Corporation, Billerica, MA), which employs the Coulter principle of impedance-based particle detection.

### Dose-response assays and combination index determination

Drug synergies were evaluated with the CompuSyn software (ComboSyn, Inc., Paramus, NJ, USA). The effectiveness of the combined drug treatments was quantified by determining the Combination Index (CI), where CI<1, CI=1, and CI>1 indicate synergism, additivity, and antagonism, respectively (Chou and Talalay, 1984).

### Colony formation assays

Single-cell suspensions of 1x10^3^ cells per well were seeded in 6-well plates and treated after 24 h with either vehicle, single drugs, or their combination. Colonies are allowed to grow for ten days and then fixed and stained with 0.5% (w/v) Crystal Violet (Sigma-Aldrich) in 20% methanol. Image acquisition with ImageJ software (public domain National Institutes of Health, USA, http://rsbweb.nih.gov/ij/) was used to score the number of colonies.

### Spheroid assays

A total of 1x10^4^ cells were seeded in 24-well ultra-low attachment culture plates (Corning, NY, USA) in the presence of the specified inhibitor(s) in serum-free media supplemented with 20 ng/ml EGF (Gibco), 20 ng/ml bFGF (Gibco), 10 µg/ml heparin (Sigma-Aldrich), 1:50 (v/v) B-27 supplement 50x (Gibco), 5 µg/ml insulin solution human (Sigma-Aldrich) and 0.5 µg/ml hydrocortisone (Sigma-Aldrich). At the end of the experiment, spheroids were incubated with 0.5 mg/ml 3-(4,5-dimethylthiazol-2-yl)-2,5-diphenyltetrazolium bromide (MTT, Sigma-Aldrich) for 2-3 h until fully stained. Plates were scanned, spheroids were scored, and spheroid area and volume were quantified by image acquisition with ImageJ software.

### Cell cycle analysis

Cells were collected by centrifugation after trypsinization, resuspended in PBS, and fixed dropwise with 70% (v/v) cold ethanol for at least 2 h. Then, cells were centrifuged, washed twice with PBS, resuspended in PBS containing 0.2 mg/ml DNAse free RNAse A (Roche, Basel, Switzerland), and incubated for 1 h at 37 °C. Prior to analysis, 50 µg/ml propidium iodide (PI) was added. Samples were analyzed using a Gallios multi-color flow cytometer instrument (Beckman Coulter, Brea, CA, USA) set up with the 3 lasers 10 colors standard configuration. Excitation was done with a blue (488 nm) laser. Forward scatter (FS), side scatter (SS), red (620/30 nm) fluorescence emitted by PI were collected. Red fluorescence from PI was projected on a 1,024-channel monoparametric histogram. Aggregates were excluded by gating single cells by their area vs. peak fluorescence signal. DNA analysis (ploidy analysis) from 10^4^ cells on single fluorescence histograms was done using MultiCycle software (Phoenix Flow Systems, San Diego, CA, USA).

### Western blot analysis

Cells were washed twice with ice-cold PBS and incubated for 30 min on ice with RIPA buffer containing 50 mM Tris (pH 8.0), 150 mM sodium chloride, 1% Triton X-100, 0.5% sodium deoxycholate, 0.1% sodium dodecyl sulfate (SDS), 1% protease inhibitor cocktail set III and 1% phosphatase inhibitor cocktail set IV (EMD Millipore Corporation). Cells were scraped, sonicated, and centrifuged at 12,000 g for 20 min at 4 °C. Supernatants were recovered, and the protein content was quantified with the BCA kit (Pierce Biotechnology).

Equal amounts of protein per sample were size-separated by electrophoresis on SDS-polyacrylamide gels and electroblotted onto Immobilon^®^-P polyvinylidene fluoride (PVDF) transfer membranes 0.45 µm pore size (Merck Millipore Ltd., Cork, IRL). After 1 h of blocking at room temperature with 5% skim milk in PBS 0.1% Tween, blots were incubated with the specific primary antibodies overnight at 4 °C. Then, membranes were treated with the appropriate horse radish peroxidase (HRP)-labeled secondary antibody for 1 h at room temperature. All blots were treated with Immobilon^®^ Western Chemiluminescent HRP Substrate (EMD Millipore Corporation, Burlington, MA, USA) and developed after exposure to an autoradiography film (VWR International, Radnor, PA, USA). The primary antibodies used were Phospho-Akt (#9271), mTOR (#2972), and Phospho-mTOR (#5536) from Cell Signaling (Beverly, MA, USA); GLS1 (ab93434), GLUD (ab166618) and MYC (ab32072) from Abcam (Cambridge, UK); GAC (19958-1-AP) and KGA (20170-1-AP) from Proteintech (Chicago, IL, USA); HIF1α (sc-13515) from Santa Cruz Biotechnology; and β-actin (#69100) form MP Biomedicals (Santa Ana, CA, USA). The secondary antibodies used were anti-mouse (ab6728) and anti-rabbit (ab6721) from Abcam.

### Measurement of extracellular metabolites

Media samples were collected at the beginning and the end of the incubation and frozen until analyzed. At the same time points, cell number was determined for normalization purposes. All the biochemical assays were carried out under exponential growth conditions. Glucose, lactate, glutamate, and glutamine concentrations from cell culture media were determined using a COBAS Mira Plus spectrophotometer (HORIBA ABX Diagnostics, Kyoto, Japan) to monitor the production of NAD(P)H in specific reactions for each metabolite at 340 nm wavelength. Glucose concentration was measured using hexokinase and glucose-6-phosphate dehydrogenase coupled enzymatic reactions (ABX Pentra Glucose HK CP, HORIBA ABX, Montpellier, France). Lactate concentration was determined at 37 °C with 2.35 mg/ml NAD^+^ (Roche) in 0.15 M hydrazine 10.9 mM EDTA pH 9 buffer and 80 U/ml lactate dehydrogenase (Roche) in 51 mM ammonium sulfate pH 6.5 buffer. Glutamate concentration was assessed at 37 °C with 2.08 mM ADP, 3.35 mM NAD^+^, and 16 U/ml of glutamate dehydrogenase (Roche) in 0.28 M glycine / 0.35 M hydrazine / 1.12 mM EDTA pH 9 buffer. Glutamine concentration was calculated through its conversion to glutamate at 37 °C for 30 min with 100 mU/ml glutaminase in 0.1 M acetate pH 5 buffer and subsequently quantifying glutamate concentration as described above.

### Estimation of metabolite consumption and production rates

Net fluxes per cell of uptake and release of different metabolites (*J_met_*) were estimated from the experimentally measured variations of metabolite concentration in medium and cell number for 24 h. The estimation was performed by assuming exponential growth and constant uptake or release per cell, which corresponds to a simple model of cell growth and metabolite consumption/production:

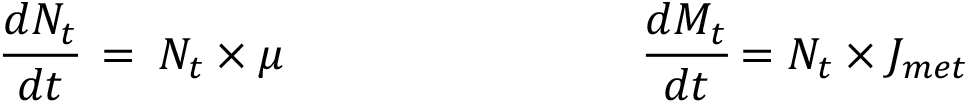

Where *N* is cell number, *M* is the quantity of metabolite, and *µ* growth rate.

### Total glutathione quantification

Fresh cells were lysed with 5% 5-sulfosalicylic acid solution, vortexed, and disrupted by two freezing/thawing cycles in liquid nitrogen and a 37 °C water bath. Cell extracts were incubated at 4 °C for 10 min and centrifuged at 10,000 g for 10 min. The reaction was initiated by mixing 150 µl of working solution (15 U/ml of glutathione reductase and 40 µg/ml of 5,5’-Dithiobis(2-nitrobenzoic acid) in 100 mM K_2_HPO_4_/KH_2_PO_4_ 1 mM EDTA pH 7.0 buffer) with 10 µl of cell extract (diluted 1:5 or 1:10) or oxidized glutathione standards (from 0 to 12.5 µM). Then, 50 µl of 0.16 mg/ml NADPH solution were added to the samples, and the increase in absorbance with time was measured at 340 nm wavelength. Total glutathione concentration was normalized by protein content.

### Mouse xenografts and in vivo drug studies

Animal experiments were carried out at the Parc Científic de Barcelona - Parc de Recerca Biomèdica de Barcelona (PCB-PRBB) Animal Facility Alliance, in compliance with current regulations (RD 53/2013; Directive 2010/63/UE; Decree 214/1997/GC, Order ECC/566/2015), to ensure that animal experiments are conducted in a humane and ethical manner. The protocol for this study was approved by the PCB’s Animal Experimentation Ethics Committee and the Animal Experimentation Commission of the Generalitat de Catalunya.

Palbociclib isethionate (S1579, SelleckChem) was resuspended at 50 mg/ml in 50 mM sodium L-lactate (Sigma-Aldrich) pH 4 buffer. Telaglenestat (S7655, SelleckChem) was dissolved at 50 mg/ml in 10 mM sodium citrate (Sigma-Aldrich) pH 2 buffer with 25% w/v (2-hydroxypropyl)-β-cyclodextrin (HP-β-CD, Sigma-Aldrich). Treatments were aliquoted into single and combined daily doses and frozen at -80 °C. HCT116^Luciferase-GFP^ cells, after a test for mycoplasma contamination, were resuspended in sterile PBS and injected (10^6^ cells in 100 µl) subcutaneously on the right flank of anesthetized 5-week-old male non-obese diabetic severe combined immunodeficiency disease (NOD-SCID) mice (strain: NOD.CB17/AlhnRj-Prkdcscid; type: Mutant congenic mouse; Janvier Labs, Le Genest-Saint-Isle, France). Tumor growth was monitored three times a week by non-invasive bioluminescence on an IVIS^®^ Spectrum In Vivo Imaging System (PerkinElmer, Waltham, MA, USA). Images and radiance were acquired 10 min after intraperitoneal injection of 150 µl of 100 mg/kg D-luciferin (BioVision Inc, Waltham, MA, USA) in sterile PBS. Eight days after tumor cells implantation, mice were randomly distributed in 4 treatment groups of fourteen animals each and administered via oral gavage with daily doses of vehicle, 100 mg/kg/day Palbociclib, 150 mg/kg/day Telaglenestat, or 100 mg/kg/day Palbociclib + 150 mg/kg/day Telaglenestat for 23 consecutive days. Mice were weighed every three days to readjust the treatment volume for precise drug dosing. Average radiance (photons (p)/s/cm^2^/steradian (sr)) was calculated for each mouse using a circular region of interest with the mouse in a prone position and normalized to the value obtained at the same area before starting treatment administrations (day 8) using the IVIS® Spectrum Living Image® 4.3.1 software (PerkinElmer). Tumor growth was also monitored thrice a week from day 15 by measuring tumor volume with a Vernier caliper using the modified ellipsoid volume formula: Tumor volume (mm^3^) = (length × width^2^) × π/6. Tumor growth inhibition (TGI) rate was calculated as TGI = (1-(V_(T,t)_/V_(T,0)_)/(V_(C,t)_/V_(C,0)_)) x 100 (Li et al., 2019), where time 0 is the first measurement of tumor volume on Day 15. At the end of the treatment, mice were euthanized, and the tumors were collected for further analysis. Animals were allowed to form tumors up to 1.5 cm in diameter, at which point they were euthanized.

### Cell line generation *ex vivo*

Three tumors for each treatment group were excised, placed in Petri dishes with DMEM/F12 medium, cut into fragments with a sterile scalpel, and subjected to digestion with 1 mg/ml collagenase type IV (Sigma-Aldrich) for 30 min at 37 °C under aseptic conditions. Then, tissue was centrifuged and digested with Trypsin EDTA solution C to generate a single cell suspension, filtered through a 40 µm cell strainer, and plated in DMEM/F12 with 2.5 mM L-glutamine, 12.5 mM D-glucose, 10% FBS, penicillin (50 U/ml), and streptomycin (50 µg/ml), obtaining twelve independent cell lines (three for each treatment condition).

### Immunohistochemistry (IHC) staining of mouse xenograft tissues

Mouse xenografts were dissected immediately after surgical resection. Three tumors for each treatment condition were fixed in 4% paraformaldehyde (PFA) and paraffin-embedded in blocks for histology assessment. Serial 4-µm tumor sections were obtained from the PFA-fixed, paraffin-embedded tumor tissues by microtome cuts. Immunohistochemistry staining was performed by the Francis Crick Institute (UK) Experimental Histopathology Laboratory on the Discovery Ultra Ventana platform (Roche). The antibody dilutions of 1:750 for CD31 (ab182981, Abcam) and of 1:1000 for KI67 (ab15580, Abcam) were used, with antigen retrieval using Cell Conditioning 1 (CC1, Roche) for 48 minutes and primary antibody incubation for 60 minutes. All slides were counterstained with Harris hematoxylin nuclear stains (Leica Biosystems, Wetzlar, Germany), dehydrated, cleared, and mounted in a Tissue-Tek Prisma® automated slide stainer (Sakura Finetek, Torrance, CA, USA). Slides were imaged on the Zeiss Axio Scan.Z1 slide scanner (Carl Zeiss Microscopy GmbH, Jena, Germany), and the percentage of KI67 and CD31 positive cells was evaluated using QuPath version 0.4.2 (Bankhead et al., 2017). A board-certified pathologist (A.S-B.) assessed the mitotic index by counting mitoses in 10 high power fields (HPF, 400×) on hematoxylin and eosin (H&E)-stained tumor sections.

### RNA sequencing

Total RNA was isolated using the RNeasy mini kit (Qiagen, Hilden, Germany) following the manufacturer’s instructions and including a DNase (Qiagen) digestion step. The size, concentration, and integrity of total RNA were determined by electrophoresis in a 4200 TapeStation system (Agilent, Santa Clara, CA, USA). Whole transcriptome sequencing (RNA-Seq) was performed at the Genomics Unit of CNIC (National Centre for Cardiovascular Research, Madrid, Spain). Barcoded RNA-Seq libraries were prepared with 200 ng of total RNA with RIN > 8 using the NEBNext® Ultra™ RNA Library Prep Kit for Illumina (New England Biolabs, Ipswich, MA, USA). Poly A+ RNA was purified using poly-T oligo-attached magnetic beads followed by fragmentation and first and second cDNA strand synthesis. The second strand was synthesized with uracil instead of thymine. Then, cDNA 3′-ends were adenylated, adapters were ligated, uracils were excised, and the libraries were amplified by PCR. The size of the libraries was determined using a 2100 Bioanalyzer DNA 1000 chip (Agilent), and their concentration was calculated using the Qubit® fluorometer (Life Technologies). All samples were indexed, and multiplex sequencing was conducted on a HiSeq2500 instrument (Illumina, San Diego, CA, USA) to generate 60-base reads in single-end format. FastQ files for each sample were obtained using CASAVA v1.8 software (Illumina). Reads were aligned to the Ensembl reference genome and converted to reads per gene with the STAR software.

### Gene expression analysis

The DESeq2 package for R was applied to normalize gene counts and identify differentially expressed genes across the study conditions (Love et al., 2014). Gene Set Enrichment Analysis (GSEA) was also used to infer enriched gene signatures from the Molecular Signatures Database (MSigDB) (Subramanian et al., 2005). The normalized enrichment score (NES) was used to rank the enriched gene sets in each phenotype. A false discovery rate q-value (FDR q-value) was computed to estimate the probability that a gene set with a given NES represented a false positive finding. Over-representation analysis (ORA) was applied to determine the biological functions or processes enriched in the differentially expressed genes of each treatment condition using WebGestalt (Liao et al., 2019).

Normalized Gene expression data from PALOMA2 (GSE133394) and PALOMA3 (GSE128500) was obtained from the GEO repository. The R package ssGSEA2 (sample Gene Set Enrichment Analysis 2) was used to compute the gene sets enrichment in each sample (Krug et al., 2019). The following sets from the Human Molecular Signatures Database (Liberzon et al., 2011) were used: HALLMARK MTORC1 SIGNALING, HALLMARK G2M CHECKPOINT, HALLMARK FATTY ACID METABOLISM, HALLMARK HYPOXIA, HALLMARK E2F TARGETS, HALLMARK GLYCOLYSIS, HALLMARK OXIDATIVE PHOSPHORYLATION, HALLMARK IL2 STAT5 SIGNALING, HALLMARK PI3K AKT MTOR SIGNALING, HALLMARK MYC TARGETS V1 and HALLMARK MYC TARGETS V2. The cor.test function from R was used to evaluate the Pearson Correlation between *PRMT* genes expressed in PALOMA2/3 samples (*CARM1* and *PRMT1*) and pathway enrichment scores and expression levels of Palbociclib-resistance genes (Zañudo et al., 2022; Zhu et al., 2022).

### Targeted metabolomics

Intracellular metabolites were extracted from tumors frozen in dry-ice cooled isopentane with 85:15 EtOH:PBS buffer and then sonicated 3 times for 5 seconds each, submerged in liquid nitrogen for 30 seconds, and thawed at 95 °C twice. Then, extracts were centrifuged at 20,000 g for 5 minutes at 4 °C, supernatants were collected, and protein content was determined. Cell media were collected at the beginning and the end of a 24-hour incubation in exponential growth, and cell number was determined for normalization purposes. Extracts from tumors and media were analyzed using the Absolute IDQ p180 kit (Biocrates Life Sciences AG, Innsbruck, Austria). Briefly, samples, calibration standards, and quality controls were dried under nitrogen gas flow at room temperature for 30 minutes. Next, 50 µl of 5% (v/v) phenylisothiocyanate (PITC) solution in 1:1:1 ethanol:water:pyridine solvent were added, incubated for 20 min at room temperature, and dried under nitrogen gas flow for 1 hour. Then, the metabolites were resuspended in 300 µl of 5 mM ammonium acetate in methanol for 30 minutes with agitation. Samples were diluted 1:1 with Milli-Q water for tandem mass spectrometry coupled to high-performance liquid chromatography (HPLC/MS/MS) or diluted 1:10 with FIA mobile phase additive (provided by the kit) for flow injection analysis (FIA) coupled to MS/MS. Analyses were performed with an Agilent 1290 Infinity Ultra-High Performance Liquid Chromatography (UHPLC) system (Agilent, Santa Clara, CA, USA) coupled with an MS/MS Sciex Triple Quad 6500 (AB Sciex LLC, Framingham, MA, USA) using electrospray ionization (ESI) in positive mode and according to the manufacturer’s instructions. The chromatographic separation was performed on an Agilent ZORBAX Eclipse XDB C18 (Agilent, Santa Clara, CA, USA) column (3.5 μm particle size, 100 × 3.0 mm) using a gradient program at a constant flow rate of 500 µl per min. The oven temperature was maintained at 50 °C. HPLC mobile phases were 0.2% formic acid in water (solvent A) and 0.2% formic acid in acetonitrile (solvent B). The elution gradient was programmed starting at 100% of solvent A for 0.5 min, then decreasing the percentage of solvent A to 5% for 5 min, holding at 95% of solvent B for 1 min, and finally re-equilibrating the column at 100% of solvent A during 0.5 min and maintaining these final conditions for 2.5 min. On the other hand, the FIA mobile phase was methanol with FIA Mobile phase Additive provided by the kit. FIA gradient started at a flow rate of 30 µl per min for 1.6 min, then increased linearly until reaching 200 µl per min during 0.8 min, maintained for 0.4 min, and finally decreased to 30 µl per min in 0.2 min. Quantitation was achieved using Multiple Reaction Monitoring (MRM) scan mode.

Data analysis was performed using Analyst (AB Sciex LLC) and MetIDQ^TM^ (Biocrates Life Sciences AG) software. A total of 21 amino acids, 21 biogenic amines, 40 acylcarnitines, 90 glycerophospholipids, 15 sphingolipids, and hexose sugars were analyzed. Data were normalized by cell number for media samples and protein for tumor samples. Clustering, heatmaps, and statistical analysis were performed using MetaboAnalyst 5.0 (Pang et al., 2022).

### Cellular bioenergetics

XFe24 Extracellular Flux Analyzer (Seahorse Bioscience, North Billerica, MA, USA) was used to measure the oxygen consumption, extracellular acidification, and proton efflux rates (OCR/ECAR/PER) in media immediately surrounding adherent cells cultured in an XFe24-well microplate (Seahorse Bioscience). One day before analysis, 7x10^4^ cells were plated in a monolayer in 100 µl, adding 150 µl of complete medium 4 h later, once cells were attached, and incubated at 37 °C and 5% CO_2_ overnight. Then, growth media were replaced by basal media (unbuffered DMEM, pH 7.4; Sigma-Aldrich) with or without glucose and incubated at 37 °C for 1 h without CO_2_. The sensor cartridge was hydrated with calibration solution (Seahorse Bioscience) overnight at 37 °C and loaded with the test reagents into the Seahorse Analyzer to calibrate the sensors. Mitochondrial function and potential were analyzed by sequential injection of 1 µM oligomycin (ATP synthase inhibitor; Sigma-Aldrich), 0.6 µM Carbonyl Cyanide m-Chlorophenylhydrazone (CCCP, mitochondrial uncoupler; Sigma-Aldrich) together with 2 mM pyruvate to achieve maximal respiration, and 2 µM rotenone and antimycin A (mitochondrial complex I and III inhibitors, respectively; Sigma-Aldrich). Cells were counted at the end of the experiments to normalize the OCR, ECAR, and PER readings. Respiration, acidification, and ATP production rates were calculated following the Agilent Seahorse XF Technology instructions.

### Building condition-specific metabolic models

Condition-specific genome-scale metabolic models (GSMMs) for control and residual tumors from *in vivo* Palbociclib and Telaglenestat treatments were reconstructed from the human GSMM Recon3D (Brunk et al., 2018). Recon3D describes the metabolic potential of the human genome, yet in any given condition, only a subset of enzymes is expressed. Hence, we reconstructed condition-specific GSMMs containing only enzymes expressed in the study conditions. More in detail, enzymes with fragments per kilobase of exon per million mapped fragments (FPKM) under 1 in all conditions were removed, provided that their removal still enabled the models to produce 50% of optimal biomass as well as the synthesis or uptake of all amino acids and biogenic amines detected with the metabolomics assays. Additionally, enzymes that were expressed below 1 FPKM in a given treatment but not in control were also removed from the treatment-specific model if the difference in gene expression to the control was statistically significant (FDR adjusted P-value <0.05).

### Quadratic multiomic metabolic transformation algorithm (qM^2^TA)

The quadratic multiomic metabolic transformation algorithm (qM^2^TA) models a metabolic transition between two states by maximizing the consistency between gene expression and metabolomics data and reaction fluxes (Figure 5A, Supplementary Figure 7). It allows the integration of both transcriptomics and metabolomics fold changes weighted by their statistical significance across both states and can identify key targets to revert the metabolic transformation.

To run qM^2^TA, we first computed the flux distribution in the control tumors condition (𝑣^/,0^, reference flux distribution) by applying the GIME3 algorithm (Schmidt et al., 2013). Briefly, this algorithm consists of a flux minimization weighted by gene expression subject to achieving 95% of the maximum biomass production and producing all measured metabolites. Next, flux variability analysis (Gudmundsson and Thiele, 2010) is used to identify the solution space within 99% of the GIME3 optimal solution. Finally, the resulting solution space is sampled using the Artificial Centering Hit-and-Run algorithm implemented into COBRApy (Ebrahim et al., 2013; Heirendt et al., 2019). The average of these flux samples is used as the control/reference flux distribution.

Then, qM^2^TA optimization is used to simulate the metabolic transition upon treatment by maximizing the consistency between gene expression and metabolite concentration fold changes and simulated reaction flux fold changes relative to the control. The optimization minimizes the difference between the resulting flux values and target flux (i.e., the product of reference flux value by fold change) for each differentially expressed metabolic gene or metabolite concentration. To give more weight to the features with more statistically significant differences, each feature (metabolite level or gene expression measure) is given a weight to the minimization inversely proportional to the P-value for the hypothesis that the fold change is different from 1. Additionally, flux variation is minimized for reactions not associated with differentially expressed genes or metabolites. The reference flux distribution scales both terms of the optimization to prevent a bias towards reactions with high reference flux.

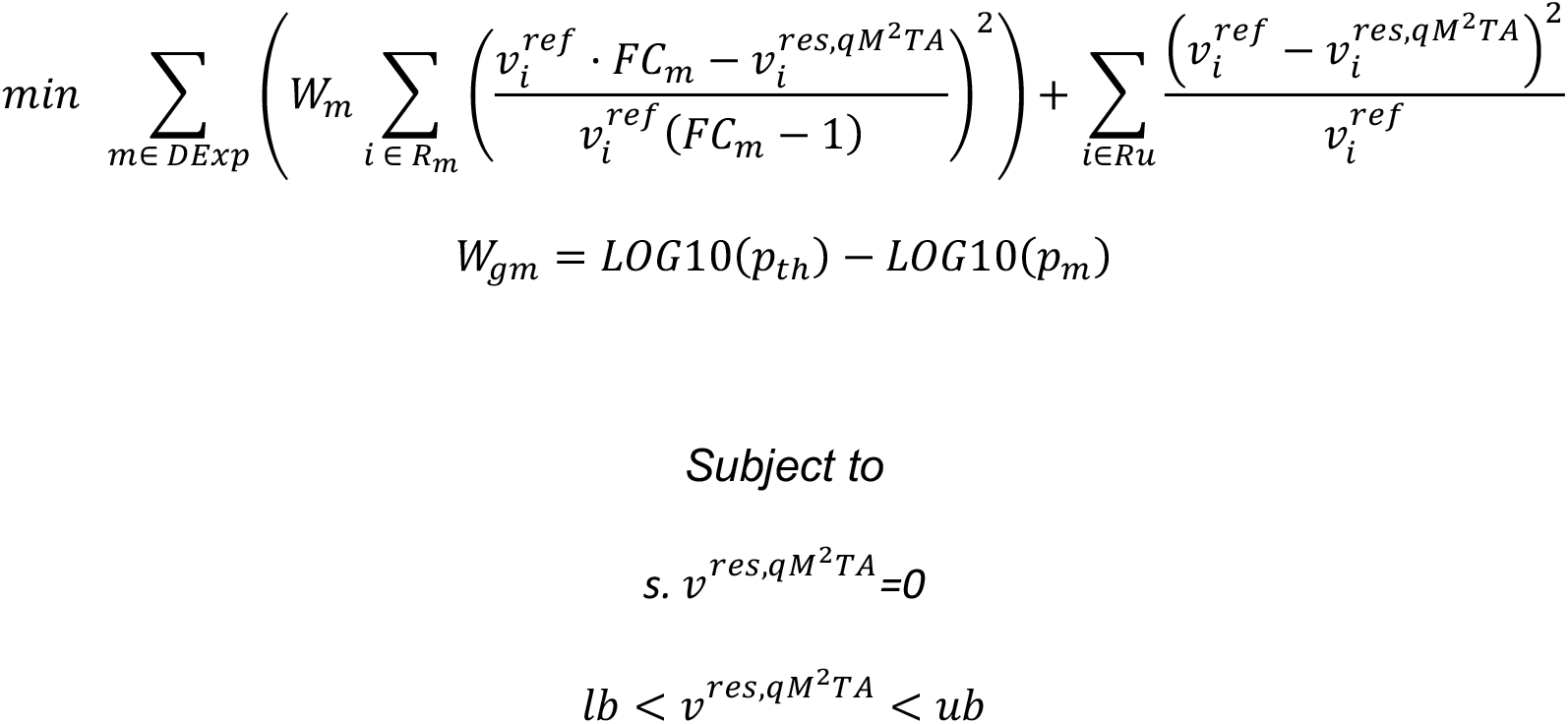

Where:

𝐷𝐸𝑥𝑝 is the list of differentially expressed genes and differentially abundant metabolites between the control and a target condition.

𝑊_𝑚_ is the weight given to measure 𝑚.

𝑅_𝑚_ are the reactions associated with measure 𝑚. For genes, this information is taken from the gene-reaction rules defined in Recon3D. Metabolomics measures are mapped to a sink reaction consuming the measured metabolite.

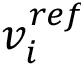 is the reference flux value (i.e., computed for the control condition) for reaction 𝑖.

𝐹𝐶*_m_* is the gene expression or metabolite concentration fold change relative to the control.

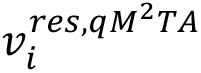 is the simulated flux value for reaction *i* in the condition of interest.

𝑅𝑢 are the reactions not associated with any differentially expressed genes.

𝑠 is the stoichiometric matrix of the condition-specific GSMMs. The product of such matrix with 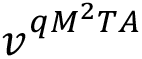 is set to 0 to define the steady-state constraint (i.e., inputs and outputs fluxes for each metabolite must be balanced).

*lb* and *ub* define reaction flux lower and upper bounds, respectively. For reactions identified as inactive when building condition-specific GSMMs, both *lb* and *ub* will be 0, preventing the reaction from carrying any flux. To simulate the effect of the glutaminase inhibitor, in the samples dosed with Telaglenestat or with the combination of Palbociclib and Telaglenestat, the maximum flux through glutaminase reaction (*ub*) was constrained to a maximum value of half the reference flux 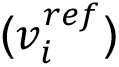.

𝑊_g_ is the weight given to each measured gene or metabolite.

𝑝_th_ is the P-value threshold used to define a fold change in gene expression or metabolite levels as differentially expressed. The threshold was defined as 0.25 FDR-adjusted P-value.

𝑝*_m_* is the FDR-adjusted P-value for measure 𝑚 for the fold change between the condition under study and the control.

Comparing the flux values simulated in a given treatment 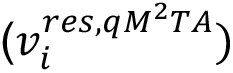 to those in control 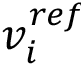 can yield a unique insight into the metabolic reprogramming underlying the adaptation to the treatment-induced stress. Reactions were grouped into metabolic pathways to facilitate interpreting the results, and the fold changes of total flux values through each pathway relative to the control were computed. Reactions were assigned to metabolic pathways based on KEGG annotations for the genes catalyzing them.

Similar to MTA and rMTA (Valcarcel et al., 2019; Yizhak et al., 2013), qM^2^TA can identify gene targets that can disrupt a particular metabolic transformation, in this case, the metabolic adaptation to Palbociclib or Telaglenestat. The potential of each metabolic gene to impede such metabolic transformation is evaluated by systematically running qM^2^TA while reducing the maximum flux (*ub*) of reactions catalyzed by a given gene to 50% of the flux value in the control condition 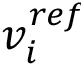. This enables testing each gene’s capacity to facilitate or disrupt the metabolic transformations associated with Palbociclib or Telaglenestat. This is complemented by testing the capacity of each gene knockdown (KD) to switch from the drug-adapted state to the control state (Valcarcel et al., 2019). The latter is achieved by running the minimization of metabolic adjustment (MOMA) algorithm while constraining the maximum flux through the reactions of a given gene to 50% of the flux value computed in the drug-adapted state 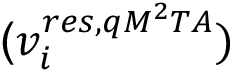. The MOMA algorithm simulates the effect of perturbations by seeking the flux distribution that minimizes the variation in reaction fluxes compared to *v^res^* and subject to the constraints of the gene KD (Segre et al., 2002).

To evaluate the potential of each target, a transformation score (TS) (Valcarcel et al., 2019; Yizhak et al., 2013) is computed as follows:

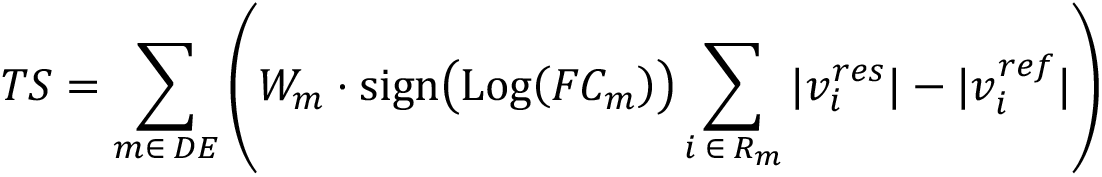

Where,

𝑇𝑆 is the transformation score.

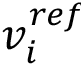 is the resulting flux distribution after either running qM^2^TA with no genes KD (wild type), running qM^2^TA with a gene KD, or running MOMA with gene KD.

A promising gene target would be any gene whose KD impairs the metabolic transformation from the control to the drug-adapted state while partially reversing the drug-adapted state towards the control state (Supplementary Figure 7). This is measured with the difference between the base TS (i.e., computed when running qM^2^TA in the wild type) and the TS when running qM^2^TA and MOMA with a gene KD.

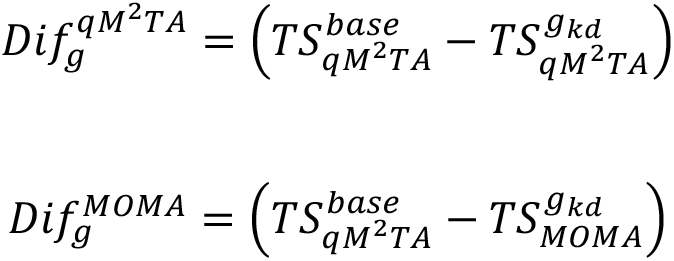

Where:

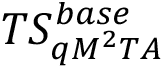 is the TS score when running qM^2^TA in the wild type (i.e., without gene KD).

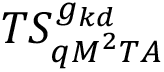 is the TS score when running qM^2^TA with gene *g* KD.

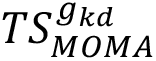 is the TS score when simulating the KD of target genes with MOMA starting from 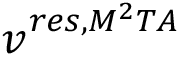

Then, the target score (𝑆g) for each gene is computed as follows:

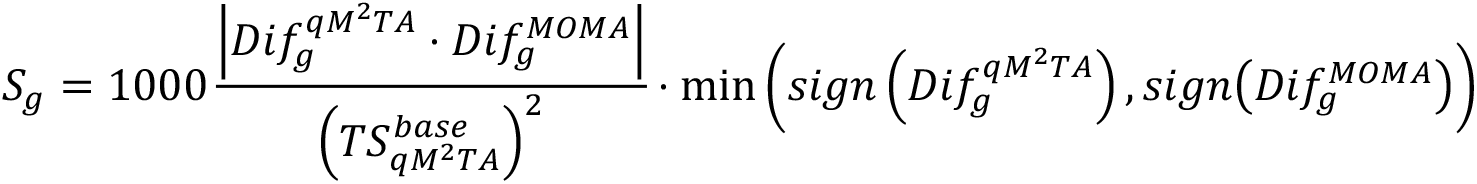

## Data availability

RNA-Seq data were deposited into the Gene Expression Omnibus database under accession number GSE245085 and are available at the following URL: https://www.ncbi.nlm.nih.gov/geo/query/acc.cgi?acc=GSE245085. All other data generated in this study are available upon request from the corresponding authors.

## Statistical analysis

Statistical analysis was conducted using GraphPad Prism 7 software (GraphPad Software, San Diego, CA, USA). All data are expressed as mean ± standard deviation (SD) unless otherwise indicated. Shapiro-Wilk test was used to assess the normal distribution of experimental data, Levene’s test was employed to evaluate the homogeneity of variances, and Dixon’s Q-test was applied to identify outliers. Kruskal-Wallis test or one-way analysis of variance (ANOVA) were used for multiple comparisons between groups. Fisher’s least significant difference (LSD) test was used to identify the groups that significantly differed from each other. Two-tailed independent sample Student’s *t*-tests were used for comparisons between two conditions, and significant differences were indicated with asterisks. Differences were considered to be significant at p < 0.05 (*), p < 0.01 (**), and p < 0.001 (***). When specified, non-significant differences (p > 0.05) are shown as n.s. Differences between groups are indicated with different letters at p < 0.05. Groups with the same letter are not significantly different (p>0.05). Concerning gene expression analysis, the DESeq2 package for R was used to evaluate the differential expression between conditions using the Wald Test. The resulting P-values were corrected for multiple testing using the Benjamini-Hochberg function built into DESeq2.

## Figure captions

**Supplementary Figure 1.**
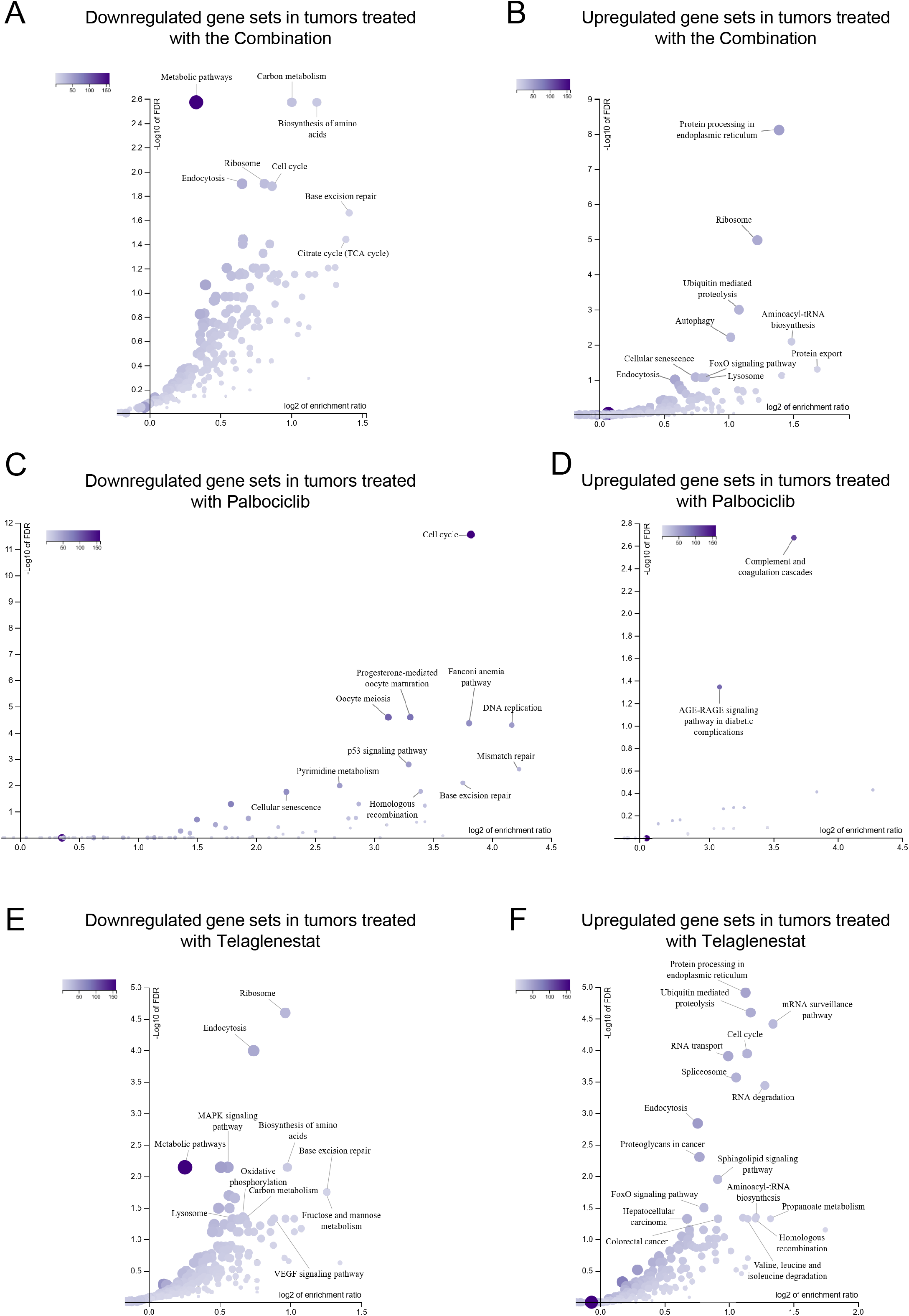
(related to Figure 1). A. Left, the chemical structure of palbociclib (PD0332991). Right, the chemical structure of Telaglenestat (CB-839). B. Synergistic antiproliferative effect of Palbociclib and Telaglenestat combined treatment. HCT116 cells were treated for 96 h at the indicated concentrations (µM) of inhibitors. The combination index (CI) results obtained with CompuSyn software (ComboSyn, Inc., Paramus, NJ, USA) revealed a strong synergy (CI<0.3) in the antiproliferative effects of Palbociclib and Telaglenestat at each dose combination tested. C. Representative IncuCyte live-cell images in phase-contrast of HCT116 cells treated with Palbociclib, Telaglenestat, or their combination at 24, 48, 72, and 96 h. Cell confluence was monitored every 3 h. Scale bar = 400 μm. D-F. Cell proliferation assay. SW403 (D), HT29 (E), and SW620 (F) cells were treated with constant-ratio increasing concentrations of Palbociclib, Telaglenestat, or their combination for 96 h, and cell quantification was assessed by Hoechst 33342 staining. Results are shown as the percentage of proliferation relative to untreated cells (mean ± SD for n=6). Top, concentration-response curves generated by data fitting with a four-parameter equation, and associated combination index (CI) values table. Bottom, dose-response cell proliferation matrix, and dose-response synergy matrix. The CI values obtained with CompuSyn software depicted a synergistic antiproliferative effect (CI<0.9) at each dose combination tested.

**Supplementary Figure 2.**
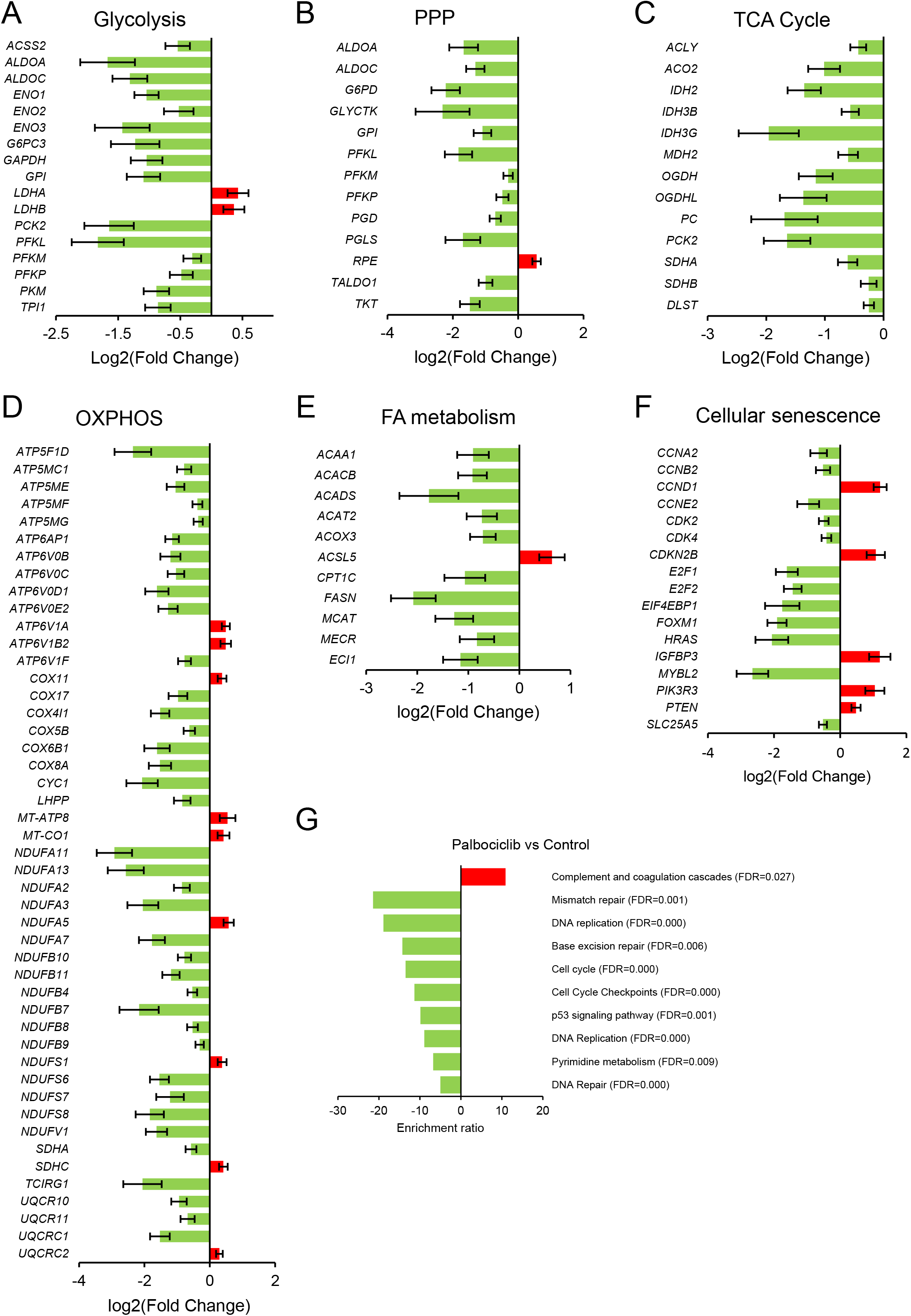
(related to Figure 2). A. Schematic outline of the experimental procedure for testing the combined therapy *in vivo*: NOD-SCID mice were injected subcutaneously with HCT116 cells (1 × 10^6^ cells/mouse) (Day 0). After seven days of tumor expansion, the mice were randomized into four groups and treated daily for 23 days with vehicle (Control), Palbociclib (PD0332991), Telaglenestat (CB-839), or their combination (PD0332991+CB-839) (n=12 per group). Tumor volume was measured with a Vernier caliper at the indicated days (orange circles). On Day 31, mice were euthanized, and tumors were extracted. B. Mean mouse weights for each treatment group were measured every 2-3 days (n=12 per group). Data are presented as mean ± SD. Yellow shadow delimits the growth curve range of male NOD.CB17-Prkdcscid/scid/Rj mice (data from Janvier Labs). C. Final body weight of individual mice at the end of the treatments (n=12 per group). Data are represented in a box and whiskers plot, with the whiskers representing the minimum and maximum values, all data points shown, and the median indicated. D. Percentage change in mouse weights from the start of the experiment normalized to the weight of Day 0. Data are shown as mean ± SD. E. Percentage change in individual mice weights at the end of the treatments (n=12 per group). Data are represented as mean ± SD. F. Representative bioluminescence imaging of mice with luciferase-expressing HCT116 subcutaneous colorectal cancer xenografts after D-luciferin injection. Tumor progression and treatment effects were monitored by 2D bioluminescence imaging (BLI) of average radiance (photons (p)/s/cm^2^/steradian (sr)). G. Representative histopathology images of Hematoxylin and Eosin (H & E) stained human colon carcinoma xenografts generated from HCT116 cells; magnification x500, scale bar = 20 μm. Circles indicate mitotic figures. H-J. Pearson linear correlation between %CD31 and %KI67 positive cells (G), mitotic count and %KI67 positive cells (H), and mitotic count and %CD31 positive cells (I). Data information: Significance was determined by ANOVA and Tukey’s multiple comparisons test with α=0.05. Statistically significant differences between conditions are indicated with different letters.

**Supplementary Figure 3.**
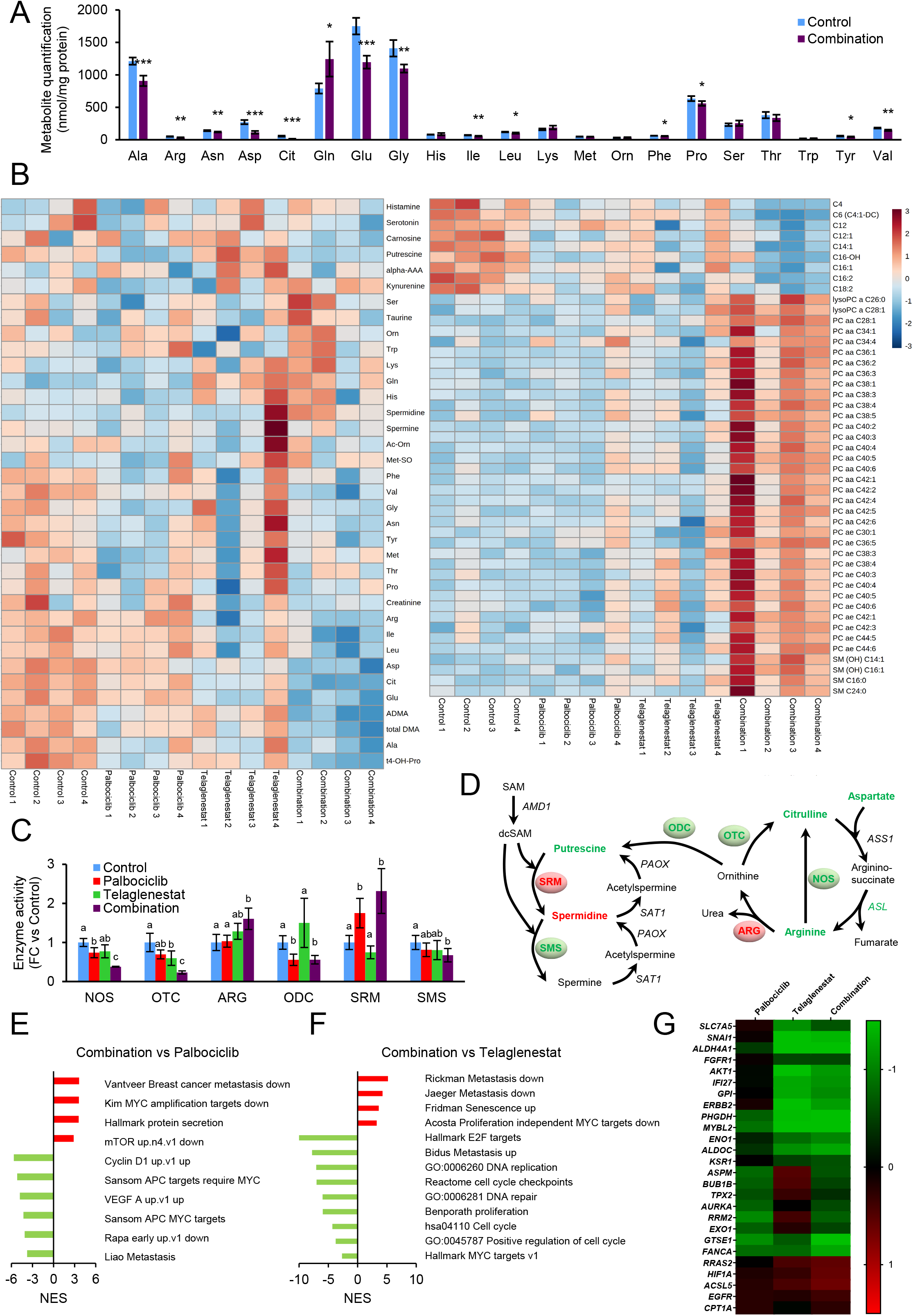
(related to Figure 3). A. Volcano plots illustrating the differential gene expression between Palbociclib, Telaglenestat, or the combined treatment and control tumors. The x-axis corresponds to the magnitude of change (log2 of the fold change), and the y-axis is the statistical significance (-log10 of the false discovery rate adjusted P-value (FDR)). Genes with FDR < 0.05 were considered statistically significant. B. Volcano plots depicting the metabolic genes differentially expressed between Palbociclib, Telaglenestat, or the combined treatment and control tumors. Metabolic genes are defined as genes that are associated with reactions or transport processes in the Recon3D genome-scale reconstruction of human metabolism. C. Top-ranked biological processes affected by Palbociclib, Telaglenestat, or the combined treatments obtained by the transcriptome profiling of HCT116 tumors. The number of differentially expressed (FDR < 0.05) genes assigned to each process is reported next to the bars. D. Volcano plots portraying the metabolic genes differentially expressed between Palbociclib or Telaglenestat and the combined treatment.

**Supplementary Figure 4.**
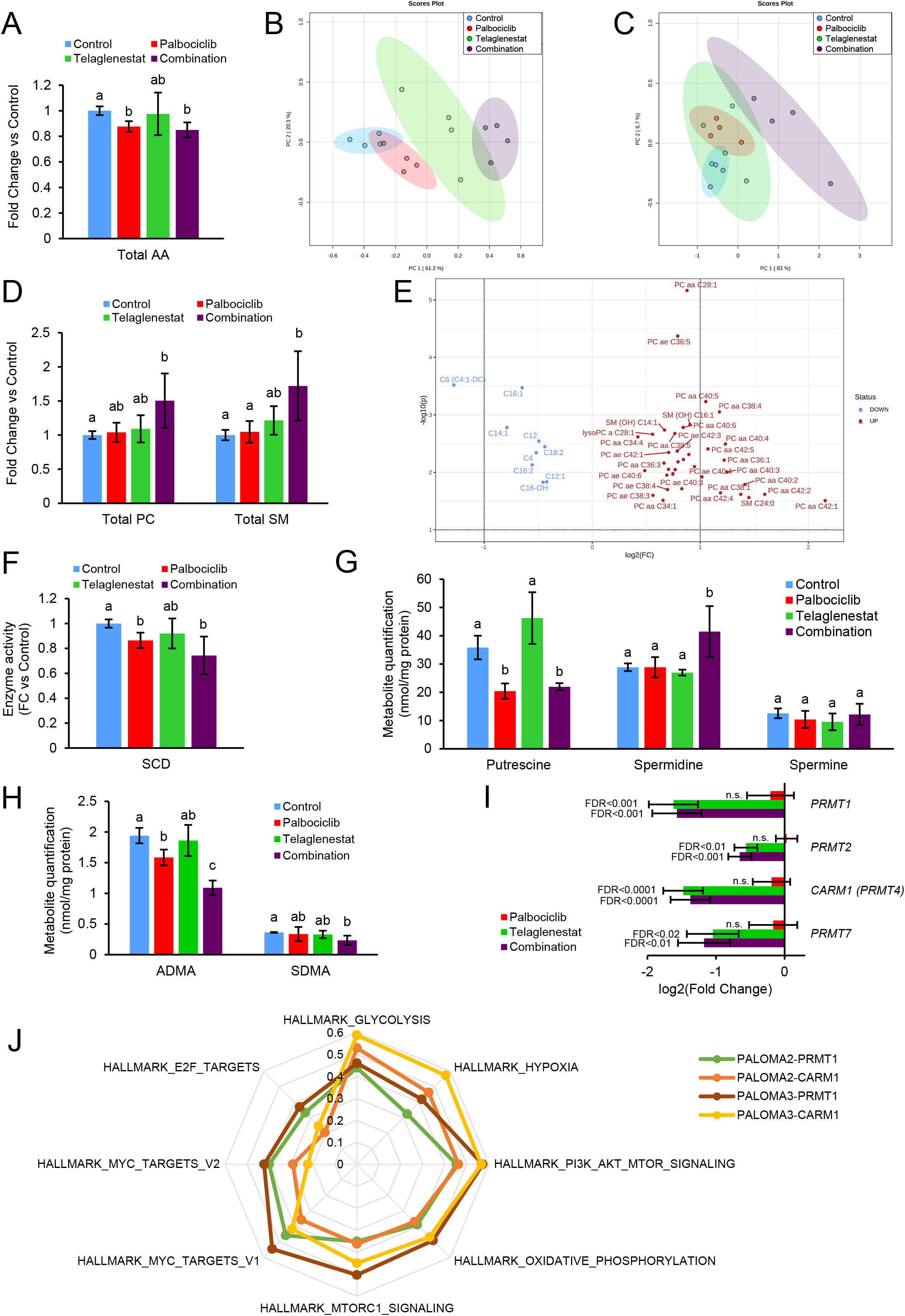
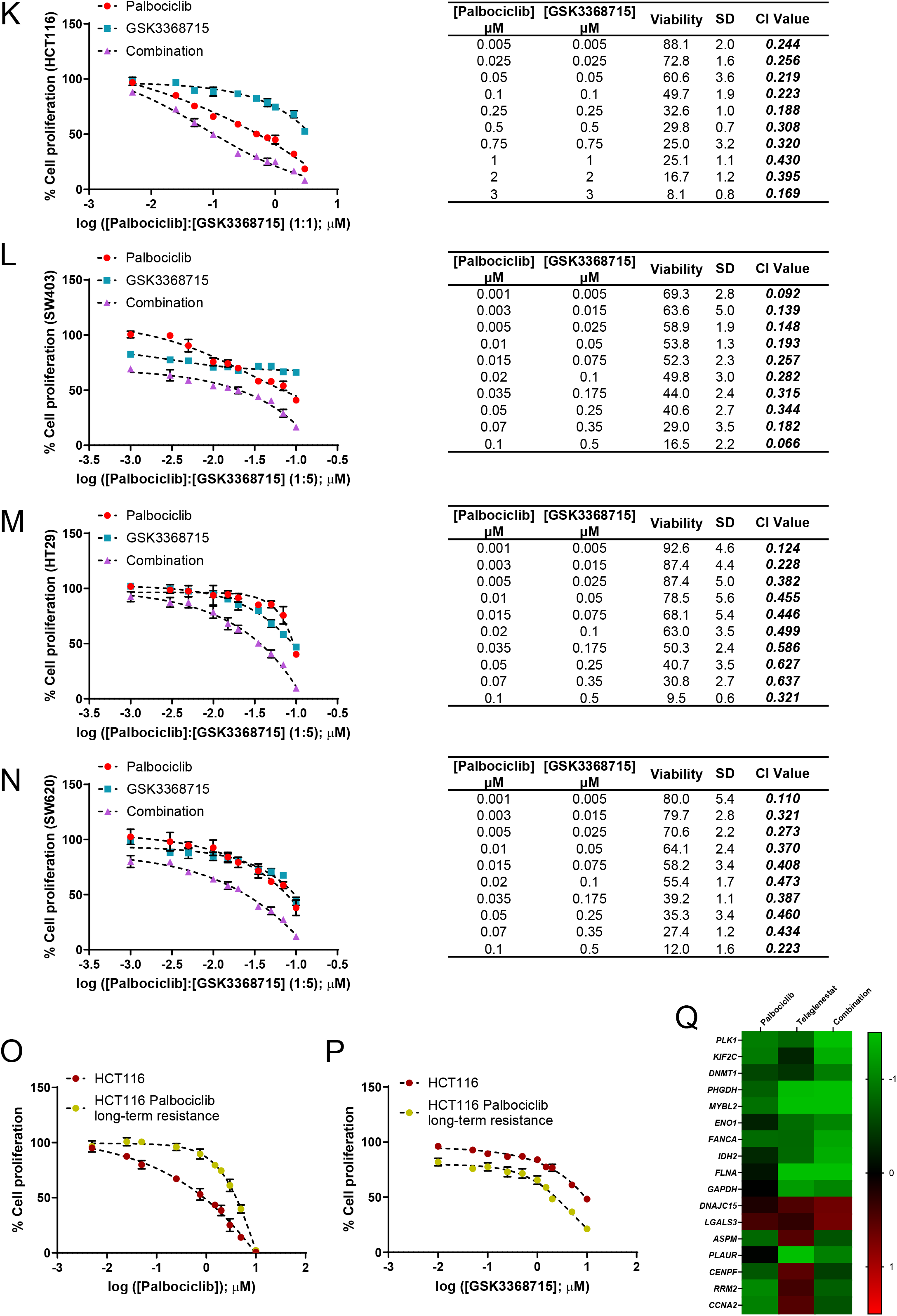
(related to. Figure 3**). Volcano plots of over-representation analysis (ORA) results.** Volcano plots representing the -log10 of FDR against enrichment ratio or NES for all the categories in the search database. Significant categories are near the upper corners. The size and color intensity of the dots are proportional to the number of overlapping genes between the gene set of the category and the differentially expressed genes of each treatment. A. Downregulated gene sets in tumors treated with the combination of Palbociclib and Telaglenestat. B. Upregulated gene sets in tumors treated with the combination of Palbociclib and Telaglenestat. C. Downregulated gene sets in tumors treated with Palbociclib. D. Upregulated gene sets in tumors treated with Palbociclib. E. Downregulated gene sets in tumors treated with Telaglenestat. F. Upregulated gene sets in tumors treated with Telaglenestat.

**Supplementary Figure 5.**
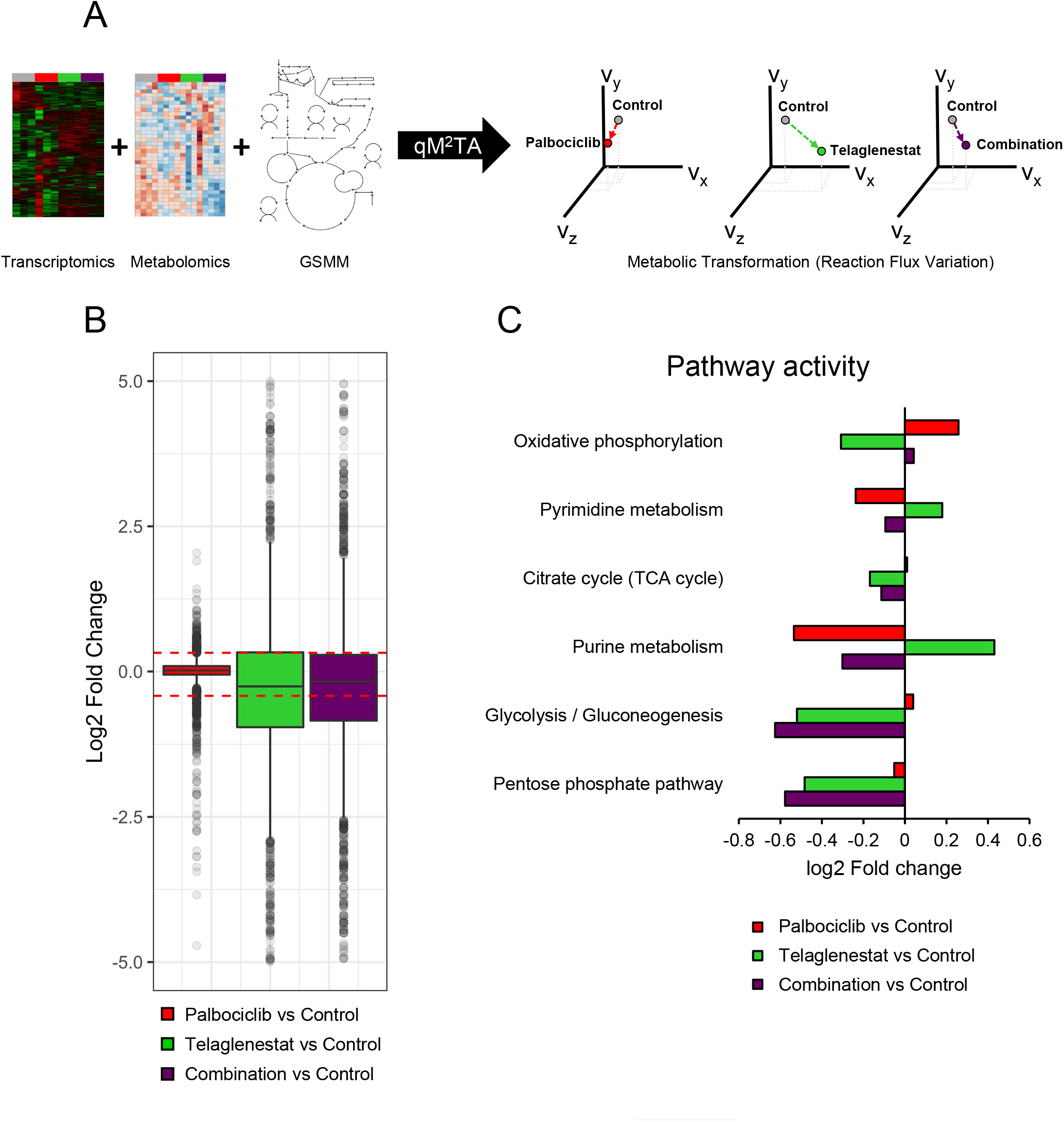
(related to Figure 3). Transcriptomic analysis of Palbociclib, Telaglenestat, and their combination treatments *in vivo*. A-F. Genes related to glycolysis (A), pentose phosphate pathway (PPP) (B), tricarboxylic acid (TCA) cycle (C), oxidative phosphorylation (OXPHOS) (D), fatty acid (FA) metabolism (E), and cellular senescence (F) that are differentially expressed (*p* adjusted < 0.05) in HCT116 tumors treated with the combination of Palbociclib and Telaglenestat compared to control tumors. G. Over-representation analysis (ORA) of downregulated (green) or upregulated (red) gene sets in tumors treated with palbociclib for 23 days (FDR < 0.05).

**Supplementary Figure 6.**
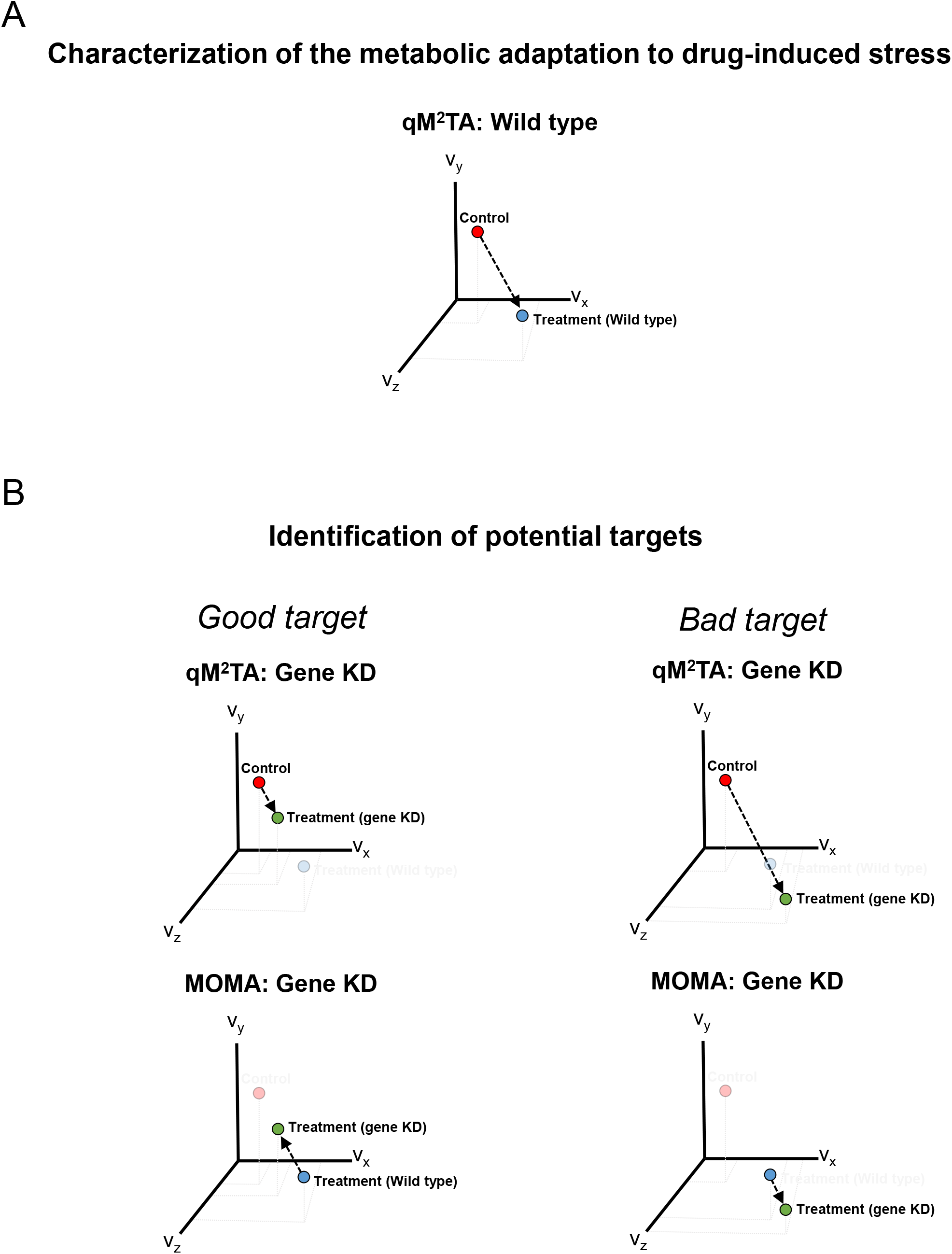
(related to. Figure 4**). Tumor metabolic profiling.** The metabolic quantification of tumors treated with vehicle, Palbociclib, Telaglenestat, or the combination of Palbociclib and Telaglenestat was determined by tandem mass spectrometry coupled to liquid chromatography (LC/MS/MS) or flow injection analysis (FIA/MS/MS). A. Quantification of total amino acids (AA) in tumors treated with Palbociclib, Telaglenestat, or the combination of Palbociclib and Telaglenestat relative to control tumors. B. Principal component analysis of the amino acid metabolic profiling of the tumors treated with vehicle, Palbociclib, Telaglenestat, or the combination of Palbociclib and Telaglenestat. C. Principal component analysis of the lipidomic profiling of the tumors treated with vehicle, Palbociclib, Telaglenestat, or the combination of Palbociclib and Telaglenestat. D. Quantification of total phosphatidylcholines (PC) and sphingomyelins (SM) in tumors treated with Palbociclib, Telaglenestat, or the combination of Palbociclib and Telaglenestat relative to control tumors. E. Volcano plot (-log10(P-value) vs log2(FC)) of the significant (P-value < 0.05) changes in lipid metabolites between tumors treated with the combination of Palbociclib and Telaglenestat and control tumors. For phosphatidylcholines (PC), the total number of carbon atoms and double bonds of the diacyl (aa) or acyl–alkyl (ae) groups is represented by Cx:y, where x indicates the number of carbons and y the number of double bonds. The same notation is used for describing the length and the number of double bonds in the acyl chain of acylcarnitines (C), lysophosphatidylcholines (lysoPC), sphingomyelins (SM) and hydroxylated sphingomyelins (SM (OH)). F. Estimation of stearoyl CoA-desaturase (SCD) enzyme activity relative to the control condition. G. Quantification of putrescine, spermidine, and spermine in tumors treated with vehicle, Palbociclib, Telaglenestat, or the combination of Palbociclib and Telaglenestat. H. Quantification asymmetric dimethylarginine (ADMA) and symmetric dimethylarginine (SDMA) in tumors treated with vehicle, Palbociclib, Telaglenestat, or the combination of Palbociclib and Telaglenestat. I. Protein arginine methyltransferases (PRMTs) genes differentially expressed (*p* adjusted < 0.05) in tumors treated with Palbociclib, Telaglenestat, or the combination of Palbociclib and Telaglenestat compared to control tumors. J. Pearson’s correlation between the enrichment of hallmark gene sets associated with Palbociclib resistance and the gene expression of *PRMT1* and *CARM1* in PALOMA-2/3 clinical trials. K-N. Cell proliferation assay for the combination of Palbociclib and the PRMT type I inhibitor GSK3368715. HCT116 (K), SW403 (L), HT29 (M), and SW620 (N) cells were treated with constant-ratio increasing concentrations of Palbociclib, GSK3368715, or their combination for 96 h, and cell quantification was assessed by Hoechst 33342 staining. Results are shown as the percentage of proliferation relative to untreated cells (mean ± SD for n=6). Left, concentration-response curves generated by data fitting with a four-parameter equation. Right, associated combination index (CI) values tables. The CI values obtained with CompuSyn software depicted a synergistic antiproliferative effect (CI<0.7) at each dose combination tested. O-P. Cell proliferation assay with Palbocilib (O) or the PRMT type I inhibitor GSK3368715 (P) in control HCT116 cells and HCT116 cells with long-term Palbociclib resistance, which were established by routinely culturing HCT116 cells in the presence of Palbociclib for four months. Cells were treated with constant-ratio increasing concentrations of Palbociclib or GSK3368715 for 96 h and cell quantification was assessed by Hoechst 33342 staining. Results are shown as the percentage of proliferation relative to untreated cells (mean ± SD for n=6). Q. Heatmap of the expression of genes associated with Palbociclib resistance, as identified in the PEARL clinical trial, in tumors treated with Palbociclib, Telaglenestat, or the combination of Palbociclib and Telaglenestat. Only the genes that are differentially expressed (p adjusted < 0.05) in HCT116 tumors treated with the combination of Palbociclib and Telaglenestat compared to control tumors are shown. Data information: Significance was determined by one-way ANOVA and Tukey’s multiple comparisons test with α=0.05. Statistically significant differences between conditions are indicated with different letters (except where stated otherwise). Shown values are mean ± SD for n=4.

**Supplementary Figure 7.**
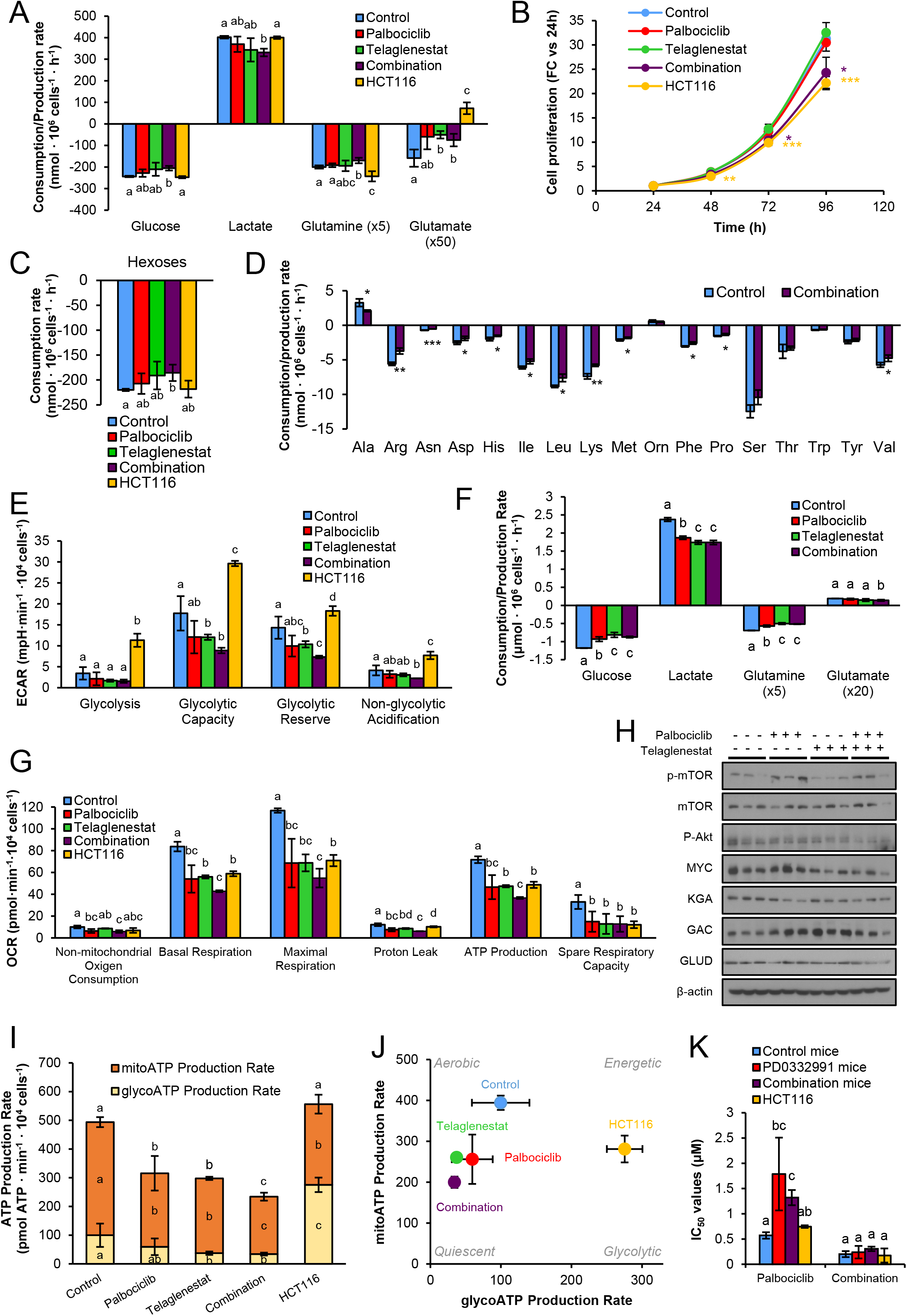
(related to Figure 5). Quadratic Metabolic transformation Algorithm (qM^2^TA). A. qM^2^TA applied to the characterization of the metabolic adaptation to drug-induced stress. qM^2^TA is used to simulate the flux changes associated with adaptation to drug-induced stress starting from the control. A hypothetical transition is shown in a 3D space for the flux through the reactions (v_x_, v_y_, v_z_). B. Identification of potential targets. Targets are evaluated by simulating the capacity of gene knockdowns (KD) to inhibit the metabolic transformation (qM^2^TA: Gene KD) and reverting the drug-adapted metabolic state (MOMA: Gene KD). A good target would be one that partially impedes the metabolic transformation from control to drug-induced stress while also allowing to partially revert the drug-induced stress metabolic state to the control state.

**Supplementary Figure 8.**
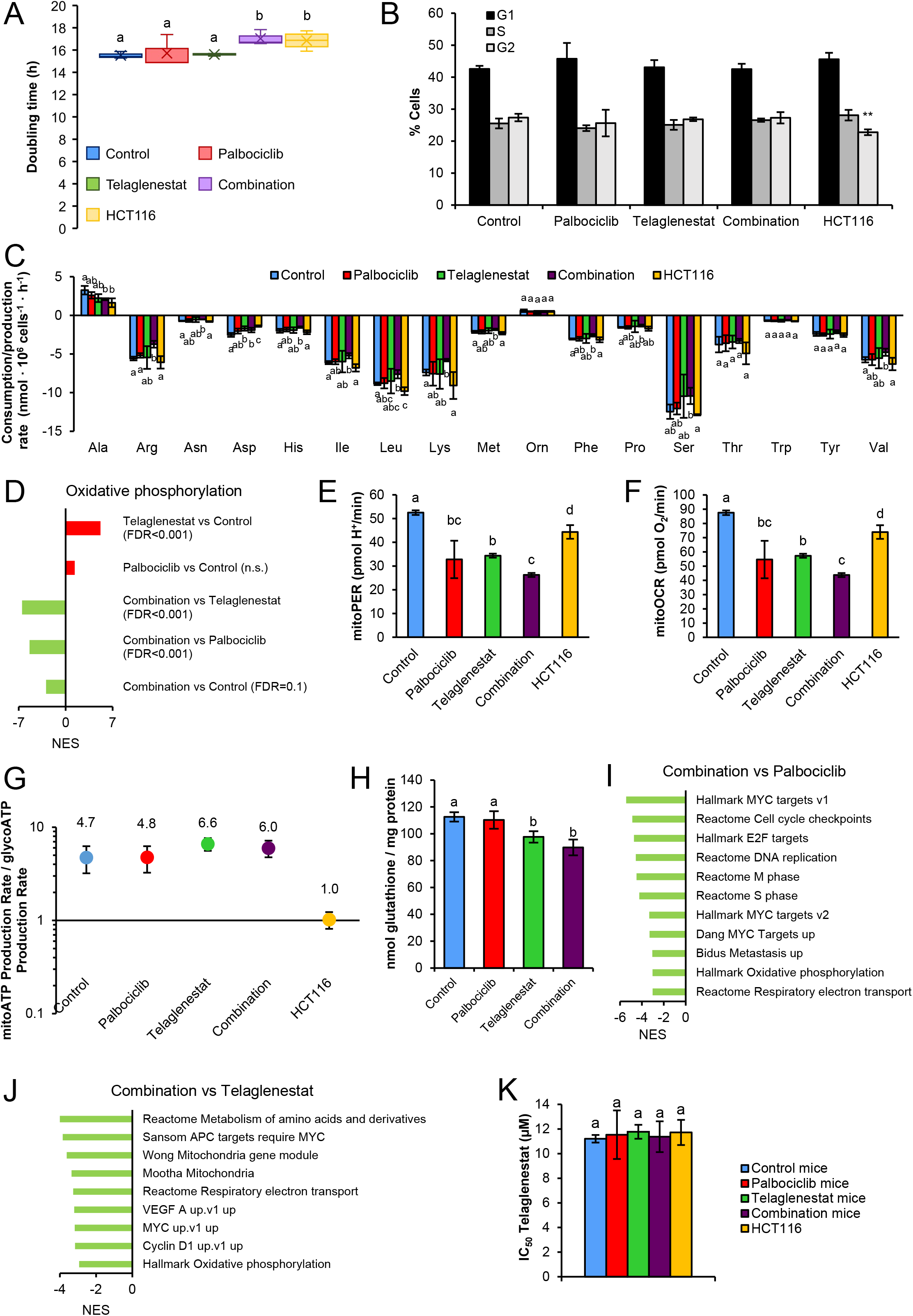
(related to. Figure 6**). Characterization of cell lines derived from HCT116 xenografts.** Twelve cell lines were obtained from HCT116 tumors from mice that had been treated daily with vehicle, Palbociclib, Telaglenestat, or the combination of Palbociclib and Telaglenestat for 23 days (three cell lines per condition). All cell lines were grown in the absence of chemotherapeutics. A. Cell doubling time of cell lines derived from tumors that had been treated with vehicle, Palbociclib, Telaglenestat, or the combination of Palbociclib and Telaglenestat, and parental HCT116 cells. B. Cell cycle distribution in cell lines derived from tumors that had been treated with vehicle, Palbociclib, Telaglenestat, or the combination of Palbociclib and Telaglenestat, and parental HCT116 cells determined by flow cytometry. C. Amino acid consumption and production rates measured by tandem mass spectrometry coupled to liquid chromatography (LC/MS/MS) after 24 h of incubation with fresh media and normalized to cell number. D. Normalized enrichment scores for the oxidative phosphorylation gene set between the cell lines derived from tumors. E. Mitochondria-derived acidification (mitoPER) assessed with a Seahorse analyzer. Data are normalized to cell number and shown as the mean of three independent cell lines for each condition ± SD with n=5. F. Mitochondrial oxygen consumption rate (mitoOCR) measured with a Seahorse analyzer. Data are normalized to cell number and shown as the mean of three independent cell lines for each condition ± SD with n=5. G. Representation of the metabolic index (or ATP rate index) as the ratio of the mitochondrial ATP (mitoATP) production rate to the glycolytic ATP (glycoATP) production rate as a quantitative metric of the cellular metabolic phenotype. H. Total glutathione quantification normalized to protein content for the cell lines derived from tumors that had been treated with vehicle, Palbociclib, Telaglenestat, or the combination of Palbociclib and Telaglenestat. I-J. Gene set enrichment analysis (GSEA) of cells obtained from tumors subjected to the combined therapy compared with cells derived from tumors treated with Palbociclib (I) or Telaglenestat (J) alone. K. IC_50_ values for Telaglenestat in cell lines derived from tumors that had been treated with vehicle, Palbociclib, Telaglenestat, and the combination of Palbociclib and Telaglenestat *in vivo*. Bars represent the mean of the IC_50_ values of three independent cell lines for each condition ± SD with n=6. Data information: Ala, alanine; Arg, arginine; Asn, asparagine; Asp, aspartate; Cit, citrulline; Gln, glutamine; Glu, glutamate; Gly, glycine; His, histidine; Ile, isoleucine; Leu, leucine; Lys, lysine; Met, methionine; Orn, ornithine; Phe, phenylalanine; Pro, proline; Ser, serine; Thr, threonine; Trp, tryptophan; Tyr, tyrosine; Val, valine. Shown values are the mean of three independent cell lines for each condition ± SD for n=3 (except otherwise indicated). Significance was determined by one-way ANOVA and Tukey’s multiple comparisons test with α=0.05. Statistically significant differences between conditions are represented with different letters.

**Supplementary Table 1 Summary statistics of the RNA-sequencing differential gene expression analysis performed *in vivo* and *ex vivo*.** The analysis was performed with the DESeq2 package for R and P-values were adjusted for multiple testing using the FDR method. Metabolic genes, defined as genes that are associated with reactions or transport processes in the Recon3D genome-scale reconstruction of human metabolism, are indicated.

**Supplementary Table 2. Correlation between the expression of *PRMT1* and *CARM1* genes and the gene sets enrichment in PI3K/AKT/mTOR signaling, OXPHOS, MYC signaling, mTOR signaling, hypoxia, glycolysis, and E2F signaling pathways in the PALOMA 2/3 clinical trials.** Gene enrichment was computed with the ssGSEA2 package for R, and Pearson correlation was evaluated with the cor.test function. The resulting P-values were adjusted for all tested pairs using the FDR method.

**Supplementary Table 3. Correlation between the expression of *PRMT1* and *CARM1* genes and genes associated with Palbociclib resistance or sensitivity in the PALOMA 2/3 clinical trials.** Pearson Correlation was evaluated with the cor.test function from R. The resulting P-values were adjusted for all tested pairs using the FDR method. Palbociclib resistance genes were identified in PALOMA-2/3 trials comparative biomarker analyses (Zañudo et al., 2022; Zhu et al., 2022).

**Supplementary Table 4. Reaction and pathway flux values computed with the quadratic metabolic transformation algorithm.** Simulated flux values in xenograft tumors for individual reactions and pathways and their variation in response to Palbociclib, Telaglenestat, or their combination are provided. Flux values were obtained by integrating transcriptomics and metabolomics with the quadratic metabolic transformation algorithm and have arbitrary units.

**Supplementary Table 5. Gene target scores computed with the quadratic metabolic transformation algorithm.** Positive gene target scores represent the putative capacity of a gene knockdown to revert the metabolic adaptation to Palbociclib or Telaglenestat. Gene targets that are significantly downregulated (adjusted P-value<0.05) at the gene expression level by either Palbociclib or Telaglenestat are indicated.

**Supplementary Table 6. The synergistic antiproliferative effect of Palbociclib and Telaglenestat combined treatment is conserved *ex vivo*.** Cells obtained from tumors from mice that have been administered with Palbociclib and Telaglenestat combination were treated for 96 h at the indicated concentrations (µM) of Palbociclib and Telaglenestat in a constant ratio (1:10). The combination index (CI) results obtained with CompuSyn software revealed a synergy (CI<1) in the antiproliferative effects of Palbociclib and Telaglenestat at each dose combination tested.

## Notes

### Competing Interest Statement

The authors have declared no competing interest.

### Summary of Updates

1. Co-correponding author affiliation. 2. New experiments and results. 3. New discussion.

