## Supplementary Tables 2, 3, 6 for "Glutaminase as a metabolic target of choice to counter acquired resistance to Palbociclib by colorectal cancer cells" (from another version of this preprint: version 2, 2024-01-11)

**Supplementary Table 1. Summary statistics of the RNA-sequencing differential gene expression analysis performed *in vivo* and *ex vivo*.** The analysis was performed with the DESeq2 package for R and P values were adjusted for multiple testing using the FDR method. Metabolic genes, defined as genes that are associated with reactions or transport processes in the Recon3D genome-scale reconstruction of human metabolism, are indicated.

**Supplementary Table 2. Correlation between the expression of *PRMT1* and *CARM1* genes and the gene sets enrichment in PI3K/AKT/mTOR signaling, OXPHOS, MYC signaling, mTOR signaling, hypoxia, glycolysis, and E2F signaling pathways in the PALOMA 2/3 clinical trials.** Gene enrichment was computed with the ssGSEA2 package for R, and Pearson correlation was evaluated with the cor.test function. The resulting P values were adjusted for all tested pairs using the FDR method.

**Supplementary Table 3. Correlation between the expression of *PRMT1* and *CARM1* genes and genes associated with Palbociclib resistance or sensitivity in the PALOMA 2/3 clinical trials.** Pearson Correlation was evaluated with the cor.test function from R. The resulting P values were adjusted for all tested pairs using the FDR method. Palbociclib resistance genes were identified in PALOMA-2/3 trials comparative biomarker analyses (Zañudo et al., 2022; Zhu et al., 2022).

**Supplementary Table 4 . Reaction and pathway flux values computed with the quadratic metabolic transformation algorithm.** Simulated flux values in xenograft tumors for individual reactions and pathways and their variation in response to Palbociclib, Telaglenestat, or their combination are provided. Flux values were obtained by integrating transcriptomics and metabolomics with the quadratic metabolic transformation algorithm and have arbitrary units.

**Supplementary Table 5. Gene target scores computed with the quadratic metabolic transformation algorithm.** Positive gene target scores represent the putative capacity of a gene knockdown to revert the metabolic adaptation to Palbociclib or Telaglenestat. Gene targets that are significantly downregulated (adjusted P value<0.05) at the gene expression level by either Palbociclib or Telaglenestat are indicated.

**Supplementary Table 6. The synergistic antiproliferative effect of Palbociclib and Telaglenestat combined treatment is conserved *ex vivo*.** Cells obtained from tumors from mice that have been administered with Palbociclib and Telaglenestat combination were treated for 96 h at the indicated concentrations (µM) of Palbociclib and Telaglenestat in a constant ratio (1:10). The combination index (CI) results obtained with CompuSyn software revealed a synergy (CI<1) in the antiproliferative effects of Palbociclib and Telaglenestat at each dose combination tested.

### Supplementary Tables 2, 3 and 6

### Supplementary Table 2

| PRMT gene | Pathway | PALOMA2 Pearson correlation | PALOMA2 Pearson FDR | PALOMA3 Pearson correlation | PALOMA3 Pearson FDR |
|---|---|---|---|---|---|
| *PRMT1* | HALLMARK_PI3K_AKT_MTOR_SIGNALING | 0.453 | 1.71E-23 | 0.573 | 1.89E-26 |
| *CARM1* | HALLMARK_PI3K_AKT_MTOR_SIGNALING | 0.462 | 2.84E-24 | 0.566 | 6.44E-26 |
| *PRMT1* | HALLMARK_OXIDATIVE_PHOSPHORYLATION | 0.389 | 4.02E-17 | 0.490 | 6.22E-19 |
| *CARM1* | HALLMARK_OXIDATIVE_PHOSPHORYLATION | 0.371 | 1.41E-15 | 0.468 | 3.44E-17 |
| *PRMT1* | HALLMARK_MYC_TARGETS_V1 | 0.458 | 6.00E-24 | 0.545 | 7.73E-24 |
| *CARM1* | HALLMARK_MYC_TARGETS_V1 | 0.358 | 1.40E-14 | 0.417 | 1.01E-13 |
| *PRMT1* | HALLMARK_MYC_TARGETS_V2 | 0.399 | 4.72E-18 | 0.422 | 5.32E-14 |
| *CARM1* | HALLMARK_MYC_TARGETS_V2 | 0.291 | 7.20E-10 | 0.224 | 1.71E-04 |
| *PRMT1* | HALLMARK_MTORC1_SIGNALING | 0.351 | 4.27E-14 | 0.504 | 5.25E-20 |
| *CARM1* | HALLMARK_MTORC1_SIGNALING | 0.363 | 6.17E-15 | 0.451 | 6.71E-16 |
| *PRMT1* | HALLMARK_HYPOXIA | 0.324 | 4.48E-12 | 0.418 | 9.46E-14 |
| *CARM1* | HALLMARK_HYPOXIA | 0.462 | 2.84E-24 | 0.572 | 1.89E-26 |
| *PRMT1* | HALLMARK_GLYCOLYSIS | 0.439 | 4.79E-22 | 0.460 | 1.33E-16 |
| *CARM1* | HALLMARK_GLYCOLYSIS | 0.529 | 1.41E-32 | 0.588 | 8.02E-28 |
| *PRMT1* | HALLMARK_E2F_TARGETS | 0.333 | 1.09E-12 | 0.369 | 8.57E-11 |
| *CARM1* | HALLMARK_E2F_TARGETS | 0.206 | 1.99E-05 | 0.246 | 3.08E-05 |

### Supplementary Table 3

| PRMT gene | PALOMA2 gene id | PALOMA2 Pearson correlation | PALOMA2 Pearson FDR | PALOMA3 Pearson correlation | PALOMA3 Pearson FDR | Type |
|---|---|---|---|---|---|---|
| *CARM1* | *ENO1* | 0.510 | 7.35E-29 | 0.561 | 1.91E-24 | Resistance genes |
| *CARM1* | *TNFRSF21* | 0.405 | 8.44E-18 | 0.523 | 7.01E-21 | Resistance genes |
| *CARM1* | *GPI* | 0.395 | 5.83E-17 | 0.450 | 2.86E-15 | Resistance genes |
| *CARM1* | *ASPM* | 0.322 | 1.96E-11 | 0.349 | 2.87E-09 | Resistance genes |
| *CARM1* | *GSR* | 0.321 | 2.41E-11 | 0.390 | 2.04E-11 | Resistance genes |
| *CARM1* | *EPS8* | 0.303 | 3.51E-10 | 0.388 | 2.37E-11 | Resistance genes |
| *CARM1* | *ALDH4A1* | 0.298 | 6.79E-10 | 0.291 | 1.20E-06 | Resistance genes |
| *CARM1* | *MTOR* | 0.283 | 6.01E-09 | 0.297 | 6.56E-07 | Resistance genes |
| *CARM1* | *AKT1* | 0.282 | 6.62E-09 | 0.309 | 2.09E-07 | Resistance genes |
| *CARM1* | *CCNE1* | 0.247 | 5.86E-07 | 0.277 | 4.06E-06 | Resistance genes |
| *CARM1* | *NCK2* | 0.244 | 8.45E-07 | 0.382 | 5.52E-11 | Resistance genes |
| *CARM1* | *ACSL5* | 0.234 | 2.34E-06 | 0.242 | 6.80E-05 | Resistance genes |
| *CARM1* | *EGFR* | 0.232 | 2.92E-06 | 0.100 | 1.32E-01 | Resistance genes |
| *CARM1* | *ERBB2* | 0.223 | 7.60E-06 | 0.395 | 1.09E-11 | Resistance genes |
| *CARM1* | *CDCA7* | 0.214 | 1.83E-05 | 0.256 | 2.44E-05 | Resistance genes |
| *CARM1* | *ALDOC* | 0.212 | 2.33E-05 | 0.265 | 1.20E-05 | Resistance genes |
| *CARM1* | *SLC15A2* | 0.160 | 1.85E-03 | 0.196 | 1.54E-03 | Resistance genes |
| *CARM1* | *FANCA* | 0.158 | 2.04E-03 | 0.157 | 1.25E-02 | Resistance genes |
| *CARM1* | *GTSE1* | 0.140 | 6.85E-03 | 0.230 | 1.75E-04 | Resistance genes |
| *CARM1* | *HIF1A* | 0.139 | 7.39E-03 | 0.124 | 5.40E-02 | Resistance genes |
| *CARM1* | *NES* | 0.128 | 1.39E-02 | 0.414 | 7.17E-13 | Resistance genes |
| *CARM1* | *EAF2* | 0.126 | 1.55E-02 | 0.187 | 2.80E-03 | Resistance genes |
| *CARM1* | *EXO1* | 0.122 | 1.95E-02 | 0.323 | 4.83E-08 | Resistance genes |
| *CARM1* | *ODC1* | 0.118 | 2.36E-02 | -0.025 | 7.39E-01 | Resistance genes |
| *CARM1* | *CDH2* | 0.112 | 3.15E-02 | 0.164 | 9.41E-03 | Resistance genes |
| *CARM1* | *SLC7A5* | 0.009 | 8.79E-01 | 0.233 | 1.38E-04 | Resistance genes |
| *CARM1* | *FMO5* | -0.112 | 3.12E-02 | -0.144 | 2.38E-02 | Sensitivity genes |
| *CARM1* | *PDZK1* | -0.120 | 2.17E-02 | -0.093 | 1.59E-01 | Sensitivity genes |
| *CARM1* | *HHAT* | -0.150 | 3.53E-03 | -0.157 | 1.29E-02 | Sensitivity genes |
| *CARM1* | *RUNDC1* | -0.158 | 2.03E-03 | -0.203 | 1.06E-03 | Sensitivity genes |
| *CARM1* | *SP1* | -0.182 | 3.40E-04 | -0.184 | 3.22E-03 | Sensitivity genes |
| *CARM1* | *IL12B* | -0.185 | 2.76E-04 | -0.361 | 7.45E-10 | Sensitivity genes |
| *CARM1* | *DNAJC17* | -0.201 | 6.28E-05 | -0.296 | 7.21E-07 | Sensitivity genes |
| *CARM1* | *PTEN* | -0.212 | 2.24E-05 | -0.349 | 3.02E-09 | Tumor suppressor |
| *CARM1* | *CYP2U1* | -0.213 | 2.08E-05 | -0.474 | 5.19E-17 | Sensitivity genes |
| *CARM1* | *NAIP* | -0.236 | 1.80E-06 | -0.343 | 5.82E-09 | Sensitivity genes |
| *CARM1* | *ATF6* | -0.299 | 6.34E-10 | -0.474 | 5.19E-17 | Sensitivity genes |
| *CARM1* | *CREB5* | -0.404 | 8.44E-18 | -0.441 | 1.22E-14 | Sensitivity genes |
| *CARM1* | *NF1* | -0.421 | 2.80E-19 | -0.577 | 4.18E-26 | Tumor suppressor |
| *PRMT1* | *ALDH4A1* | 0.389 | 1.88E-16 | 0.285 | 2.02E-06 | Resistance genes |
| *PRMT1* | *EPS8* | 0.380 | 1.13E-15 | 0.432 | 5.09E-14 | Resistance genes |
| *PRMT1* | *ERBB2* | 0.365 | 1.60E-14 | 0.501 | 3.87E-19 | Resistance genes |

| | | | | | | |
|---|---|---|---|---|---|---|
| *PRMT1* | *ENO1* | 0.363 | 2.28E-14 | 0.484 | 8.68E-18 | Resistance genes |
| *PRMT1* | *GPI* | 0.355 | 1.00E-13 | 0.488 | 4.28E-18 | Resistance genes |
| *PRMT1* | *ASPM* | 0.352 | 1.59E-13 | 0.432 | 5.10E-14 | Resistance genes |
| *PRMT1* | *AKT1* | 0.318 | 3.85E-11 | 0.415 | 6.61E-13 | Resistance genes |
| *PRMT1* | *GSR* | 0.314 | 7.16E-11 | 0.316 | 1.07E-07 | Resistance genes |
| *PRMT1* | *MTOR* | 0.312 | 8.98E-11 | 0.380 | 6.62E-11 | Resistance genes |
| *PRMT1* | *EXO1* | 0.266 | 5.42E-08 | 0.384 | 4.05E-11 | Resistance genes |
| *PRMT1* | *TNFRSF21* | 0.243 | 8.88E-07 | 0.255 | 2.73E-05 | Resistance genes |
| *PRMT1* | *FANCA* | 0.223 | 7.73E-06 | 0.219 | 3.52E-04 | Resistance genes |
| *PRMT1* | *GTSE1* | 0.209 | 3.01E-05 | 0.322 | 5.44E-08 | Resistance genes |
| *PRMT1* | *NES* | 0.194 | 1.26E-04 | 0.276 | 4.78E-06 | Resistance genes |
| *PRMT1* | *MYBL2* | 0.190 | 1.79E-04 | 0.109 | 9.66E-02 | Resistance genes |
| *PRMT1* | *RRM2* | 0.171 | 8.44E-04 | 0.198 | 1.40E-03 | Resistance genes |
| *PRMT1* | *NCK2* | 0.167 | 1.04E-03 | 0.288 | 1.47E-06 | Resistance genes |
| *PRMT1* | *ALDOC* | 0.163 | 1.43E-03 | 0.253 | 3.14E-05 | Resistance genes |
| *PRMT1* | *CCNE1* | 0.161 | 1.75E-03 | 0.247 | 4.98E-05 | Resistance genes |
| *PRMT1* | *BUB1B* | 0.158 | 2.03E-03 | 0.019 | 7.90E-01 | Resistance genes |
| *PRMT1* | *TPX2* | 0.157 | 2.14E-03 | 0.137 | 3.25E-02 | Resistance genes |
| *PRMT1* | *ESR1* | 0.153 | 2.99E-03 | 0.378 | 8.44E-11 | Resistance genes |
| *PRMT1* | *SLC15A2* | 0.131 | 1.17E-02 | 0.099 | 1.32E-01 | Resistance genes |
| *PRMT1* | *ACSL5* | 0.107 | 3.90E-02 | 0.121 | 6.13E-02 | Resistance genes |
| *PRMT1* | *SLC7A5* | 0.084 | 1.15E-01 | 0.186 | 2.93E-03 | Resistance genes |
| *PRMT1* | *PRKAR2A* | -0.107 | 4.01E-02 | 0.014 | 8.56E-01 | Sensitivity genes |
| *PRMT1* | *SP1* | -0.131 | 1.21E-02 | 0.095 | 1.49E-01 | Sensitivity genes |
| *PRMT1* | *CX3CR1* | -0.133 | 1.07E-02 | -0.189 | 2.48E-03 | Sensitivity genes |
| *PRMT1* | *DNAJC17* | -0.161 | 1.75E-03 | -0.203 | 1.02E-03 | Sensitivity genes |
| *PRMT1* | *NAIP* | -0.177 | 5.33E-04 | -0.135 | 3.57E-02 | Sensitivity genes |
| *PRMT1* | *IL12B* | -0.178 | 4.76E-04 | -0.276 | 4.71E-06 | Sensitivity genes |
| *PRMT1* | *PTEN* | -0.206 | 4.11E-05 | -0.224 | 2.62E-04 | Tumor suppressor |
| *PRMT1* | *ATF6* | -0.235 | 2.08E-06 | -0.386 | 3.11E-11 | Sensitivity genes |
| *PRMT1* | *NF1* | -0.327 | 9.57E-12 | -0.512 | 5.12E-20 | Tumor suppressor |
| *PRMT1* | *CYP2U1* | -0.347 | 3.65E-13 | -0.492 | 2.17E-18 | Sensitivity genes |
| *PRMT1* | *CREB5* | -0.377 | 1.70E-15 | -0.378 | 8.40E-11 | Sensitivity genes |

### Supplementary Table 6

| | | Cell line 1 derived from tumor 1 treated with Combination | | | Cell line 2 derived from tumor 2 treated with Combination | | | Cell line 3 derived from tumor 3 treated with Combination | | |
|---|---|---|---|---|---|---|---|---|---|---|
| **[Palbociclib] μM** | **[Telaglenestat] μM** | **Viability** | **SD** | **CI Value** | **Viability** | **SD** | **CI Value** | **Viability** | **SD** | **CI Value** |
| 0.04 | 0.4 | 80.9 | 1.5 | ***0.450*** | 76.8 | 1.0 | ***0.615*** | 78.6 | 1.9 | ***0.662*** |
| 0.08 | 0.8 | 71.0 | 2.3 | ***0.398*** | 64.7 | 5.6 | ***0.386*** | 67.0 | 2.7 | ***0.400*** |
| 0.15 | 1.5 | 59.7 | 3.3 | ***0.367*** | 55.6 | 2.0 | ***0.379*** | 57.2 | 1.3 | ***0.350*** |
| 0.3 | 3 | 52.2 | 2.0 | ***0.486*** | 48.9 | 5.2 | ***0.514*** | 48.9 | 4.8 | ***0.404*** |
| 0.5 | 5 | 50.7 | 5.4 | ***0.749*** | 47.9 | 3.0 | ***0.813*** | 48.1 | 1.1 | ***0.641*** |
| 0.65 | 6.5 | 49.0 | 2.4 | ***0.891*** | 43.9 | 2.5 | ***0.869*** | 43.5 | 0.8 | ***0.632*** |
| 0.8 | 8 | 42.8 | 1.5 | ***0.797*** | 41.0 | 5.0 | ***0.940*** | 41.8 | 3.3 | ***0.704*** |
| 0.9 | 9 | 37.9 | 3.5 | ***0.698*** | 35.0 | 2.6 | ***0.834*** | 37.7 | 3.6 | ***0.627*** |
| 1.1 | 11 | 27.0 | 1.6 | ***0.476*** | 26.3 | 1.8 | ***0.764*** | 17.7 | 1.0 | ***0.238*** |
| 1.4 | 14 | 13.5 | 2.1 | ***0.245*** | 11.1 | 2.4 | ***0.616*** | 8.6 | 1.7 | ***0.140*** |

Combination Index (CI) equation of Chou and Talalay and CompuSyn software

| CI value | Agonistic effect |
|---|---|
| <0.10 | Very strong synergism |
| 0.10–0.30 | Strong synergism |
| 0.30–0.70 | Synergism |
| 0.70–0.90 | Moderate to slight synergism |
| 0.90–1.10 | Nearly additive |
| 1.10–1.45 | Slight to moderate antagonism |
| 1.45–3.30 | Antagonism |
| >3.30 | Strong to very strong antagonism |
